# Genomic analyses provide new insights into the evolutionary history and reproduction of the Paleogene relictual *Kingdonia* (Circaeasteraceae, Ranunculales)

**DOI:** 10.1101/2022.06.07.495209

**Authors:** Yanxia Sun, Xu Zhang, Aidi Zhang, Jacob B. Landis, Huajie Zhang, Hang Sun, Qiu-Yun (Jenny) Xiang, Hengchang Wang

**Author notes:** These authors contributed equally (Yanxia Sun, Xu Zhang, Aidi Zhang). Author for correspondence: Hang Sun,; Jenny Xiang,; Hengchang Wang.

## Abstract

Asexual lineages are perceived to be short-lived on evolutionary timescales due to accumulation of deleterious mutations. Hence reports for exceptional cases of putative ‘ancient asexual’ species usually raise doubts about whether such species are recently derived or engage in some undocumented sexual reproduction. However, there have been few studies to solve the mystery. The monotypic *Kingdonia* dating to the early Eocene, contains only *K. uniflora* that has no known definitive evidence for sexual reproduction nor records for having closely related congeneric sexual species, seeming to have persisted under strict asexuality for long periods of time. In this study, we use population genomic analyses to test for reproduction mode and infer the evolutionary process and mechanisms facilitating the survival of the species. Our results indicate the presence of three differentiating genetic lineages within the species and support that asexual reproduction in *K. uniflora* indicated by high allelic heterozygosity had occurred prior to the lineage differentiation (∼0.5 mya). We also detected DNA recombination events in some populations, in line with occurrence of unseen and unevenly distributed sexual reproduction among populations. However the observation of high linkage disequilibrium, relatively high ratio of *π*_N_/*π*_S_ (nonsynonymous versus synonymous nucleotide diversity), together with high allelic heterozygosity suggest the sexual reproduction is infrequent. Furthermore, we found that genes containing SNPs with elevated *F*st values are significantly enriched in functions associated with seed development, suggesting differentiation in genes regulating seed development is likely to be the key reason of the uneven distribution of sexual reproduction in *K. uniflora*. Evidence from our study supports predominate asexual reproduction in *K. uniflora*, but unseen sexual reproduction must have played a key role to ensure the long-term survival of the lineage in general. Uneven distribution of sexual reproduction in the species may be a key factor underlying the observed genetic differentiation between populations. This study provides novel insights into the reproduction and evolution of *Kingdonia*, a relict lineage evolved in the Paleogene and known for asexual reproduction, and demonstrate the power of data from population genome sequencing in resolving long-standing evolutionary questions.

## Introduction

Sexual and asexual reproduction are the two basic methods of plant reproduction. Although sexual reproduction is the predominant mode in vascular plants, asexual reproduction occurs in many taxonomic groups (Mogie 1992). Compared to sexual reproduction, asexual reproduction is often described as “a short cut” and “cost effective”, requiring no waiting time and resources for fertilization to occur, resulting in production of more offspring in less time (Corley et al. 2001). Evolutionary theory predicts that asexual reproduction is not a successful long-term strategy. In the absence of sexual reproduction, accumulation of deleterious mutations, e.g., elevated ratio of non-synonymous (selected) to synonymous (neutral) polymorphisms (*π*_N_/*π*_S_) is expected due to relaxed purifying selection in asexual lineages (Normark and Moran 2000; Ament-Velásquez et al. 2016). One consequence is the eventual extinction of asexual lineages once the accumulation reaches a high load of deleterious mutations (Muller’s ratchet; Muller 1964; Felsenstein 1974). Hence, asexual lineages are traditionally perceived as evolutionary dead-ends (Maynard-Smith 1978; Zimmer 2009). However, a number of exceptions, the so-called ‘ancient asexual’ species, e.g., darwinulid ostracods and parthenogenetic oribatid mites, have been reported (Heethoff et al. 2007; Schoen et al. 2009; Brandt et al. 2021). These lineages were suggested to have persisted under obligate asexuality over millions of years, which violates the expectation that sexual reproduction and recombination are necessary for long-term survival (Schurko et al. 2009). A long standing hypothesis is that such ‘ancient asexual’ lineages should have special adaptive mechanisms, e.g., an efficient DNA repair system, to cope with the accumulation of deleterious mutations (e.g., Birky et al. 2005; Gladyshev and Meselson 2008). Alternatively, such lineages may not be true ‘ancient asexuals’, and several species (including the famous Bdelloid rotifers) previously believed to be long-term asexuals were indeed later shown to be either recently derived or to engage in some unseen sexual reproduction (Lunt 2008; Schurko et al. 2009; Signorovitch et al. 2015; Schwander 2016; Laine 2020; Brandt et al. 2021).

In the literature, the determination of obligate asexual reproduction in such ‘ancient asexual’ lineages has usually relied on negative evidence, such as the failure to find individuals of the opposite sex for mating or failure to detect a recent sexual ancestor (Neiman et al. 2009). Such evidence is not conclusive regarding sexual reproduction is absent due to the possibility that unseen sexual reproduction may exist, and the apparent “obligate” asexuality might reflect our inability to observe sexual reproduction, making dominantly asexual lineages appear strictly asexual (Schurko et al. 2009). Hence, assessing the true mode of reproduction in putative ‘obligate asexual’ species requires more reliable methods. In contrast to previous organismal-based methods, molecular approaches provide an effective way to distinguish obligate from facultative asexual lineages. There are a number of expected genetic consequences under obligate asexual reproduction. For example, in asexual diploid species, alleles are expected to highly divergent due to the independent accumulation of mutations in the absence of segregation and genetic exchange, the well-known Meselson effect (Birky 1996; Mark Welch and Meselson 2000). Highly divergent alleles are also expected in recently derived homoploid hybrid species due to the divergence of gene copies derived from different species (Beck et al. 2012; Jaron et al. 2021).

Correspondingly, a negative value of the *F*_IS_ index, which measures the level of within-individual heterozygosity, is expected in these lineages, as observed in obligate asexuals (Balloux et al. 2003; Ament-Velásquez et al. 2016). Another well-acknowledged signature of asexual reproduction is the generation of non-random associations between loci, i.e., linkage disequilibrium (LD), which is often used for estimating the amount of asexual reproduction. Therefore, genome-wide linkage disequilibrium is expected in obligate asexuality (de Meeûsa and Balloux 2004; Henry et al. 2012; Lovell et al. 2014). Additionally, in a strictly asexual lineage, where mutation is the only source of genetic novelty and all loci show complete linkage, the genealogy of genes and genomes of the lineage are expected to be a strictly branching tree rather than a network (Normark et al. 2003). Although some of these genetic consequences related to asexual reproduction have been assessed in animals, investigating all of these consequences in a single study has been rare, and largely lacking in plant lineages appears to be asexual only. .

*Kingdonia*, represented by a single species *K. uniflora*, is one of two monotypic genera (*Kingdonia* and *Circaeaster*) in the family Circaeasteraceae (Ranunculales; Angiosperm Phylogeny Group, 2016). The genus represents an ancient lineage estimated to have diverged from its sister *Circaeaster agrestis* Maxim. during the early Eocene based on molecular dating using the DNA sequences of three chloroplast spacers (52 mya; 95% HPD=27-75 mya), 497 single-copy genes (51.8 mya; 95% HPD=31-76 mya) and whole plastome sequences (52.2 mya; 95% HPD=26-83 mya), respectively (Ruiz-Sanchez et al. 2012; Sun et al. 2020; Zhang et al. 2020). Unique among angiosperms, both *K. uniflora* and *C. agrestis* possess an unusual dichotomous leaf venation similar to that of ferns and *Ginkgo*, indicating a reversal to an ancestral venation type of vascular plants. *Kingdonia uniflora* (diploid, 2n = 18) is a herbaceous species with a genome size of ∼ 1 Gb, endemic to alpine regions of southwest China and grows in cold and humid habitats with deep humus (Sun et al. 2020). The species has a very narrow distribution being restricted to the Qinling Mountains, Minshan Mountains and Daxue-Qionglai Mountains (Fig. 1a). Notably, *K. uniflora* is well-known to produce new “individuals” by means of rhizome rupture, which occurs on rhizomes more than three years old (Lei et al. 2000; Li et al. 2003; Supplemental Fig. S1). Although production of seeds was observed occasionally, but different from its sexual sister species *C. agrestis*, the seed embryo of *K. uniflora* can only develop to the torpedo stage (Ren et al. 1998). Efforts to germinate *K. uniflora* seeds in the natural habitat and lab had a zero rate of success (Li et al. 2004). So far, field investigations throughout the species’ range found no seedlings of the species in natural populations (Li et al. 2003; Xu 2015). Available evidence suggests that the species may have evolved without sexual reproduction for long periods of time, representing a putative ‘ancient asexual’ lineage. Here we characterize genome-wide genetic variation across nearly all known *K. uniflora* populations. Specifically, we aim to determine (1) if the species is indeed an obligate asexual lineage or cryptic sexual reproduction is present, and (2) when asexual reproduction has evolved in the species. To answer these questions and better understand the evolutionary mechanisms of the species, we conducted various analyses to (a) characterize the genetic variation and population structure, (b) examine the genomic signatures of reproductive strategies, (c) reconstruct the demographic and evolution histories, and (d) identify and annotate outlier genetic variants.

**Fig. 1.**
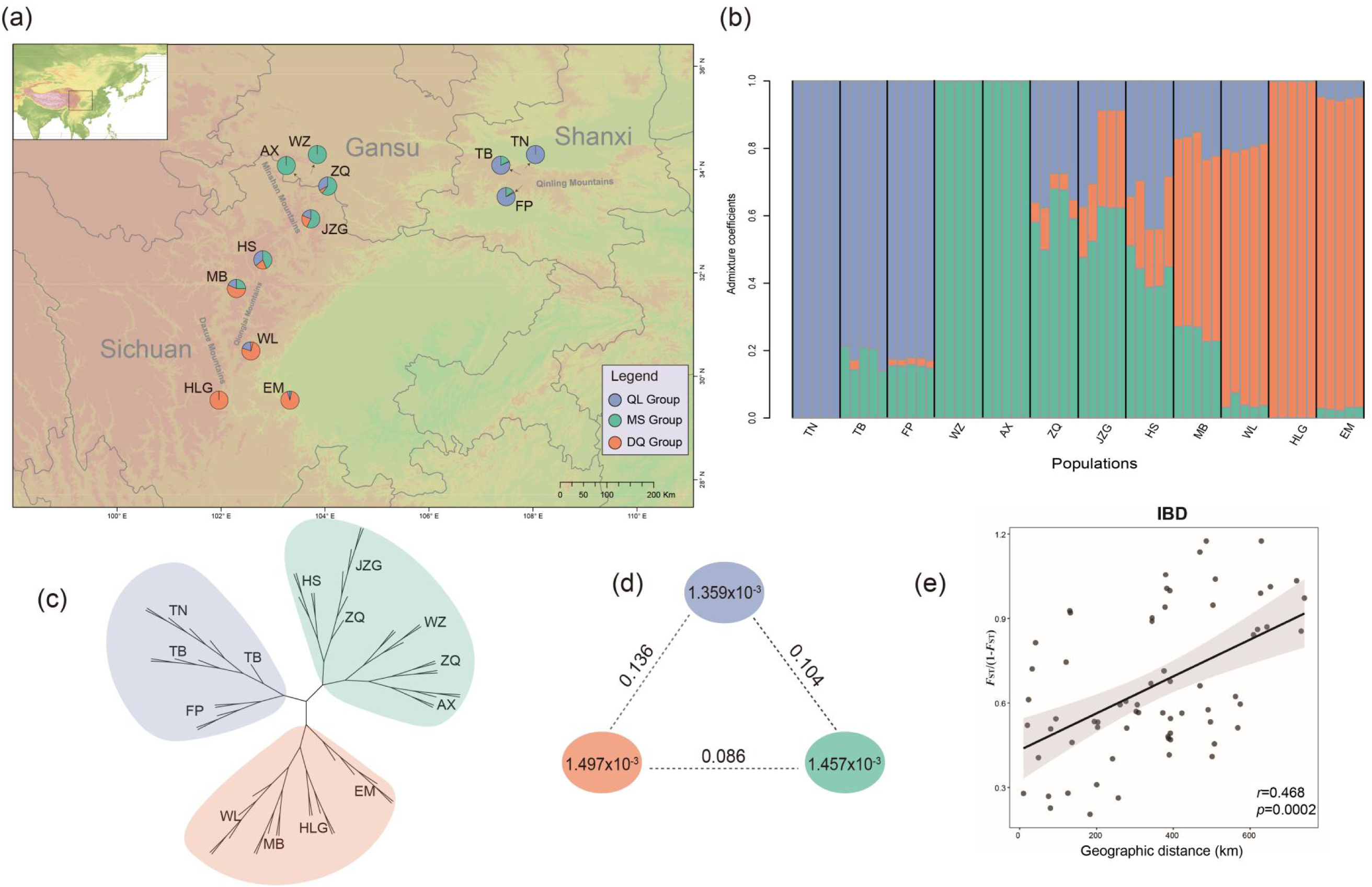
Genetic structure and diversity within *K. uniflora*. (a) Sampling sites and genetic structure detected by LEA analysis (*K* = 3 populations) mapped using ArcGIS v10.3. The genetic QL group is shown in blue, the MS group is shown in green, and the DQ group is shown in orange. (b) Genetic groups of *K. uniflora* inferred by the LEA analysis when setting *K*=3. (c) A phylogeny derived from SVDQuartets using 114,746 SNPs. (d) Nucleotide diversity (π) and population divergence (*F*_ST_) across three genetic groups. The value in each circle represents a measure of nucleotide diversity for this group, and the value on each line indicates divergence between groups. (e) Correlation between genetic distance and geographical distance (isolation by distance, IBD), as tested by Mantel test.

## Results

### Sequence data processing

We produced 1601 Gb of data containing 2,356,880,312 raw reads for 60 individuals of *K. uniflora*, and 2,342,304,596 clean reads (SRA------) after filtering. The depth of coverage for the 60 samples ranged from 21.6× (WZ4) to 34.8× (TN1), with a mean coverage of 26.6x (Supplemental Table S1). Using the *K. uniflora* reference genome (GenBank: PRJNA587615, Sun et al. 2020), we obtained 9,349,354 raw SNPs, and 8,678,779 high quality SNPs after primary standard strict filtering. Additionally, by applying deep filtering standards, we obtained 114,746 SNPs located in 960 contigs (Supplemental Table S2).

### Genetic diversity and population structure

Our LEA (Landscape and Ecological Association) analysis with the cross-entropy method failed to identify an ideal best-fit number of *K* (Supplemental Fig. S2). However, our phylogenetic analysis using the SVDQuartets method (Chifman and Kubatko 2014) revealed three clusters (supported by three quarters (361494/487635) of all the quartets), corresponding to the QL (Qingling Mountains), MS (Minshan Mountains) and DQ (Daxue-Qionglai Mountains) groups (Fig. 1c). Therefore, we repeated the LEA analysis (Fig. 1b; Supplemental Fig. S3) by setting *K*=3. The result recognized the same three groups identified by SVDQuartets. Additionally, two of the three populations from the QL group, three of the five populations from the MS group, and two of the four populations from the DQ group exhibit different levels of genetic admixture with the other groups (Fig. 1a,1b).

Nucleotide diversity (π) (the average π value) or per-site heterozygosity analysis showed that the three groups have similar genetic diversity, ranging from 1.359×10^-3^ to 1.497×10^-3^ (Table 1; Fig. 1d). The genetic differentiation statistics (fixation index; *F*_ST_) among the three groups estimated from VCFtools v0.1.16 (Danecek et al. 2011) was 0.136 between the non-adjacent QL and the DQ groups (highest), 0.086 between the adjacent MS group and DQ group (lowest), and 0.0104 between the adjacent groups MS and QL (Fig. 1d), indicating low genetic differentiation among the three groups. The Mantel test showed significant isolation by distance (IBD) (*r* = 0.468, *P* = 0.0002) (Fig. 1e). Results from the AMOVA showed small genetic variation among the three groups (5.6%; Table 2). The results also showed no significant amount of variation among individuals within populations (with a negative component value of -48%; Table 2). And most of the genetic variation in *K. uniflora* was explained by variation within individuals (with a component value of 123.83% within individuals) (Table 2).

**Table 1.**
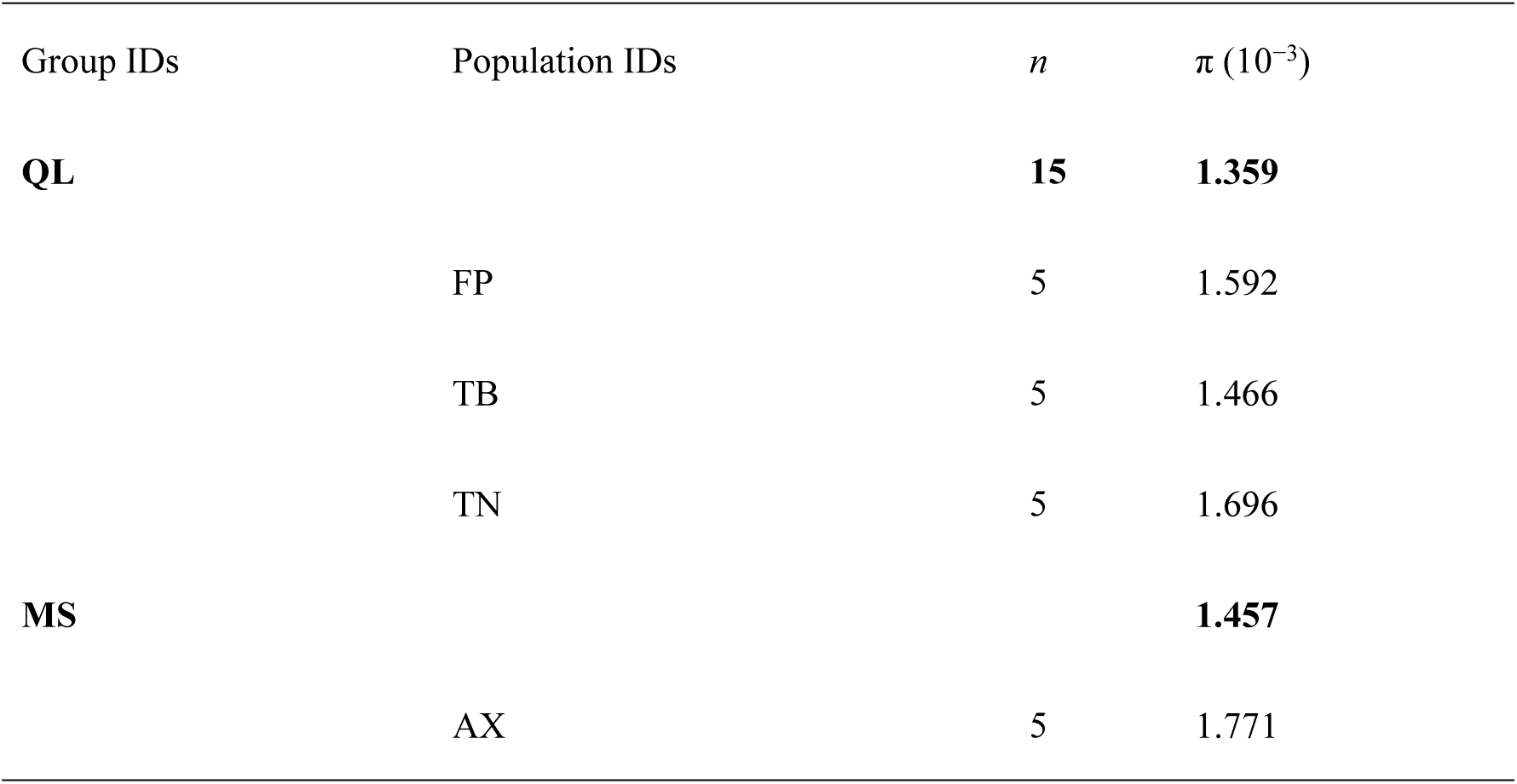
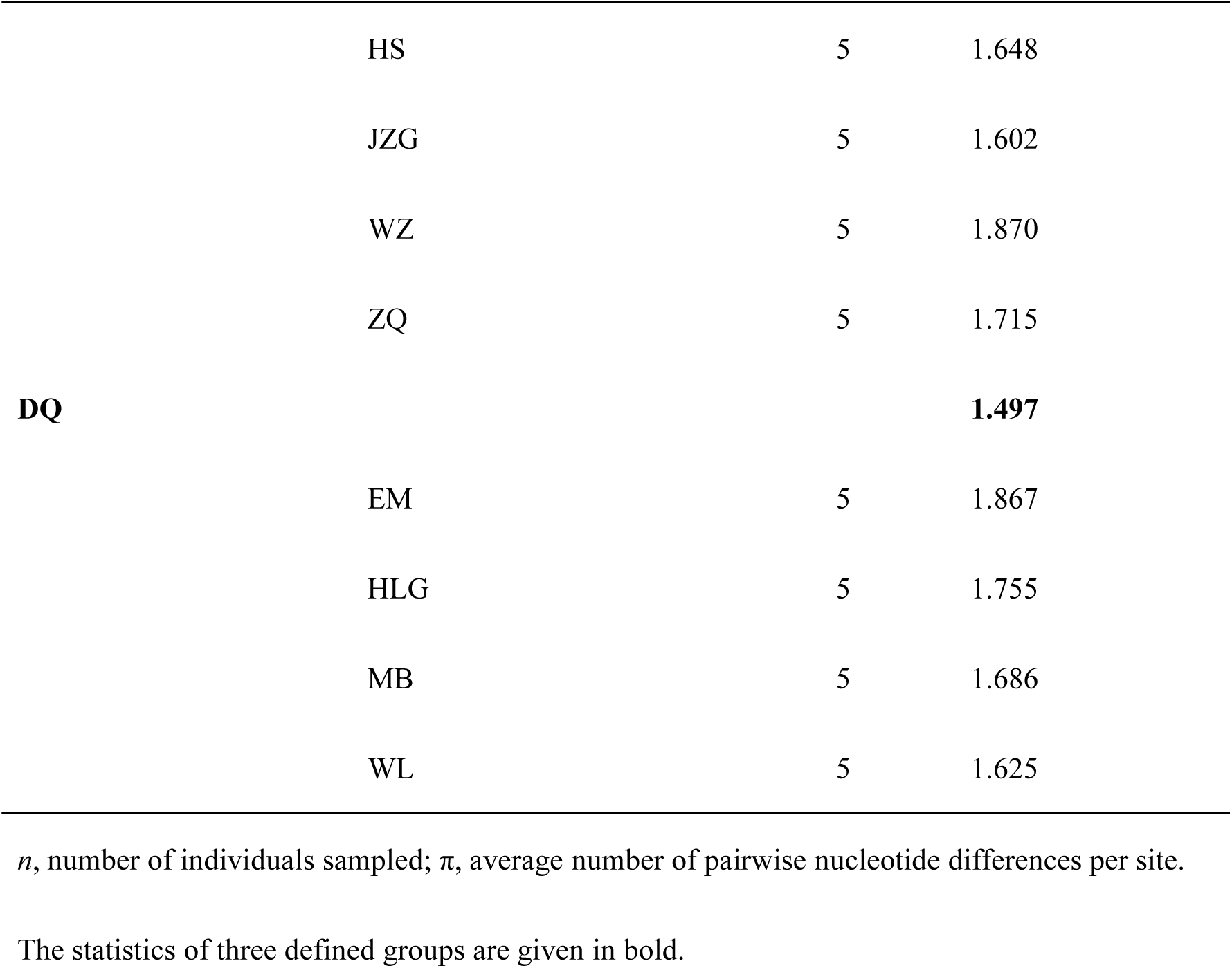
Summary of genetic diversity of K. uniflora calculated using 114,746 variant positions.

| Group IDs | Population IDs | *n* | $\pi$ ($10^{-3}$) |
| --- | --- | --- | --- |
| **QL** |  | **15** | **1.359** |
|  | FP | 5 | 1.592 |
|  | TB | 5 | 1.466 |
|  | TN | 5 | 1.696 |
| **MS** |  |  | **1.457** |
|  | AX | 5 | 1.771 |
|  | HS | 5 | 1.648 |
|  | JZG | 5 | 1.602 |
|  | WZ | 5 | 1.870 |
|  | ZQ | 5 | 1.715 |
| **DQ** |  |  | **1.497** |
|  | EM | 5 | 1.867 |
|  | HLG | 5 | 1.755 |
|  | MB | 5 | 1.686 |
|  | WL | 5 | 1.625 |

*n*, number of individuals sampled; $\pi$ , average number of pairwise nucleotide differences per site.
The statistics of three defined groups are given in bold.

**Table 2.**
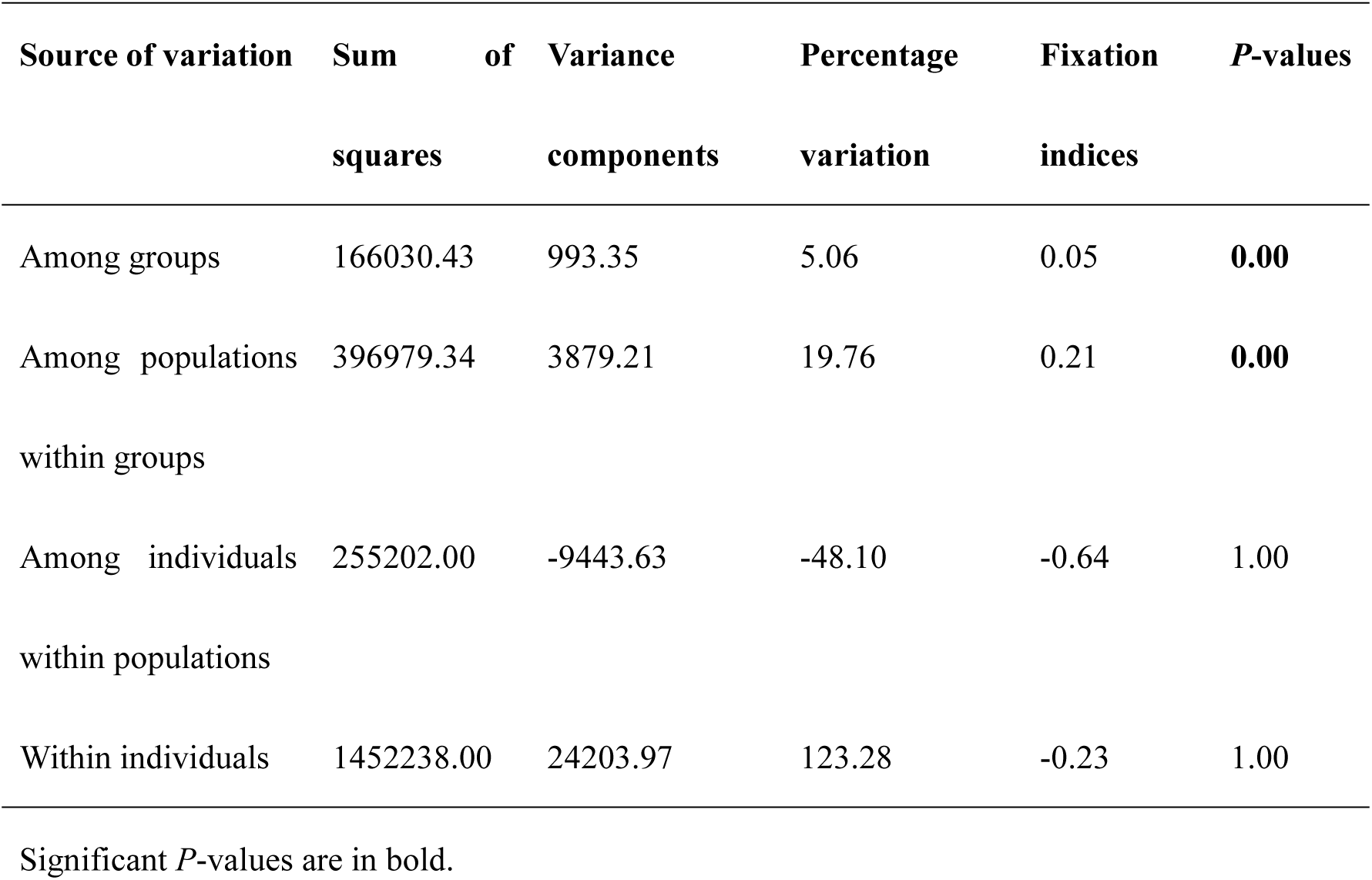
Details on the results of the analysis of the molecular variance (AMOVA).

### Genomic signatures and signs of sexual and asexual reproduction

The calculation of *F*_IS_ using VCFtools v0.1.16 (Danecek et al. 2011) revealed negative *F*_IS_ values (ranging from -0.07 to -0.47) for all 60 *K. uniflora* individuals (mean *F*_IS_=-0.26) (Fig. 2), indicating greater observed individual heterozygosity (Supplemental Table S3) than expected from random mating, a sign of asexual reproduction. The site frequency spectrum (SFS) across all 12 populations showed sites with heterozygous SNPs shared among three lineages/all 12 populations were most abundant (12,367 sites), *c.* 250 times more frequent than expected under Hardy-Weinberg equilibrium (HWE) (purple bar in Fig. 3), suggesting a large proportion of the observed excess individual heterozygosity in the 12 populations were attributed to accumulation by asexuality prior to the evolutionary divergence of the three genetic lineages (QL, MS and DQ).

**Fig. 2.**
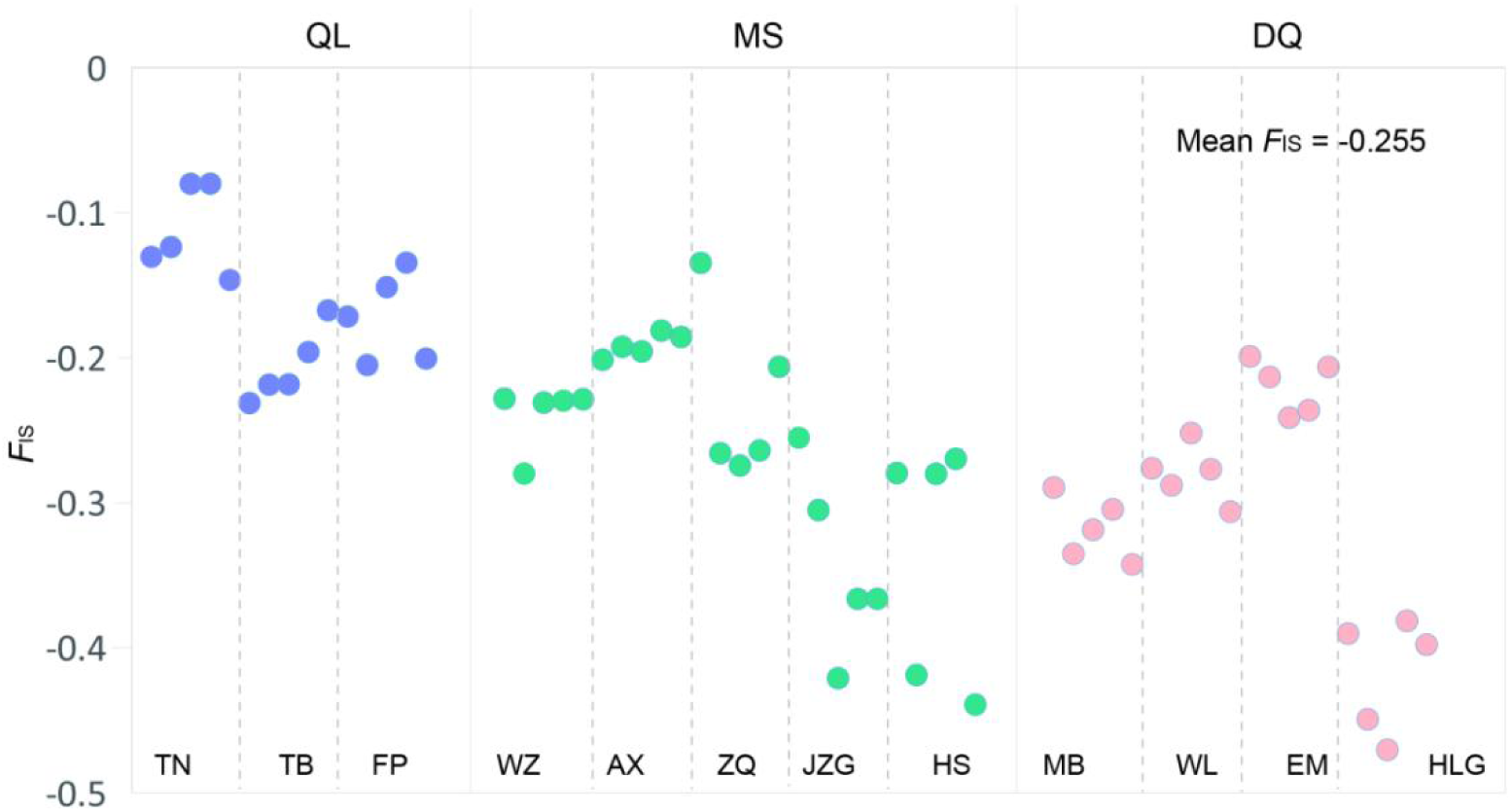
Distribution of values of inbreeding coeffificient (*F*_IS_). Under Hardy-Weinberg equilibrium *F*_IS_ is 0, negative values of *F*_IS_ indicate excess of individual heterozygosity.

**Fig. 3.**
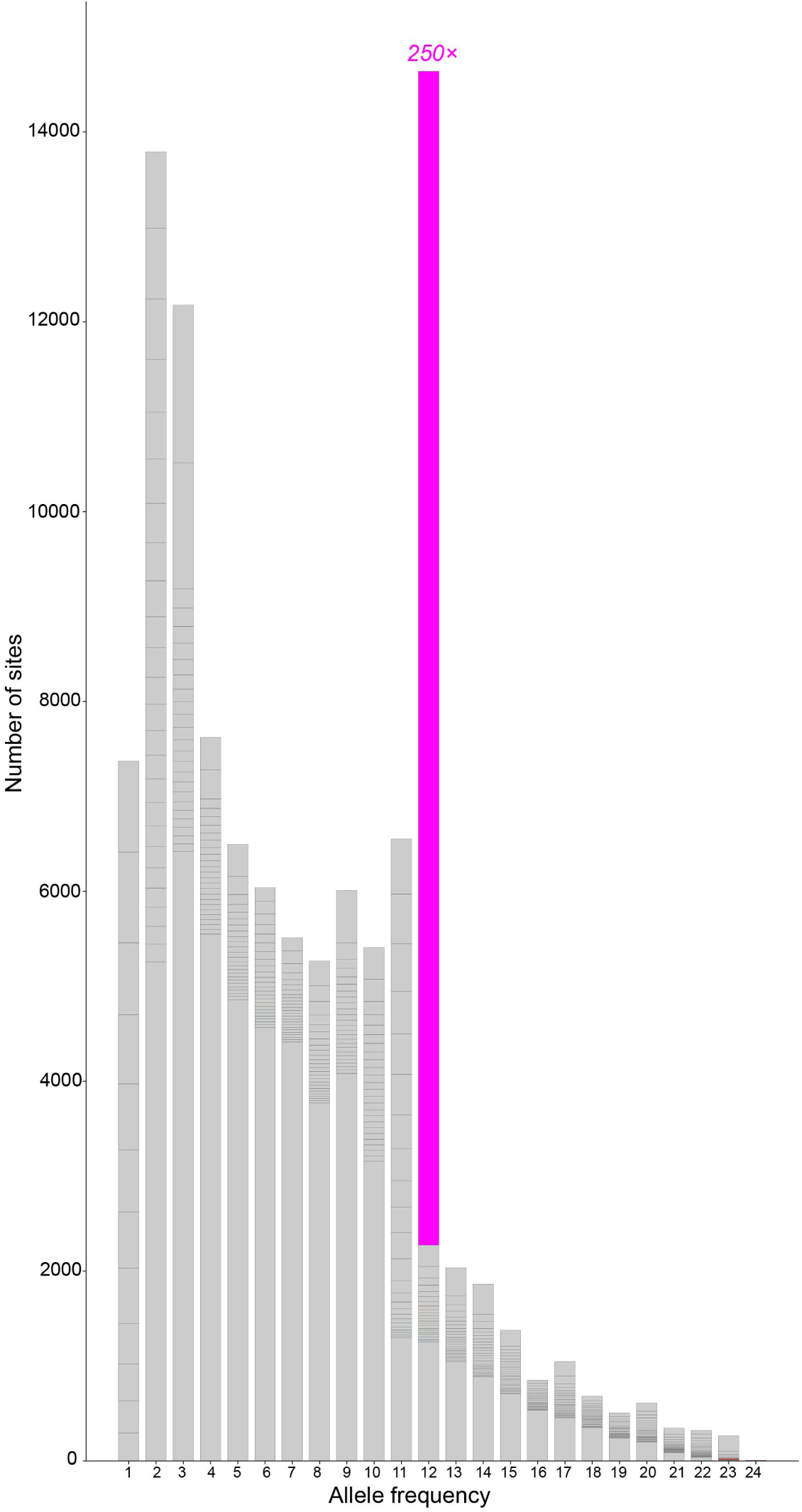
The site frequency spectrum (SFS) depicting the number of sites with different allele frequency across 12 *K. uniflora* individuals (one allele can display a maximum frequency of 24 among 12 diploid individuals). Heterozygous genotypes shared among all 12 populations are highlighted using the purple color and its excess over HWE indicated (*c*. 250 times as frequent as expected under HWE).

Result from Linkeage Disequilibrium (LD) analyses showed sites within 0-25 kb region had an average *r*^2^ of 0.60, 0.50 and 0.50, which slowly decay to *c*. 0.50, 0.40 and 0.40 after a physical distance of 300 kb in QL, MS and DQ, respectively (Fig. 4), implying very slow LD decay over very large distance. In addition, the LD analysis between pairs of SNPs up to 1000 kb obtained similar results (Fig. S4).

**Fig. 4.**
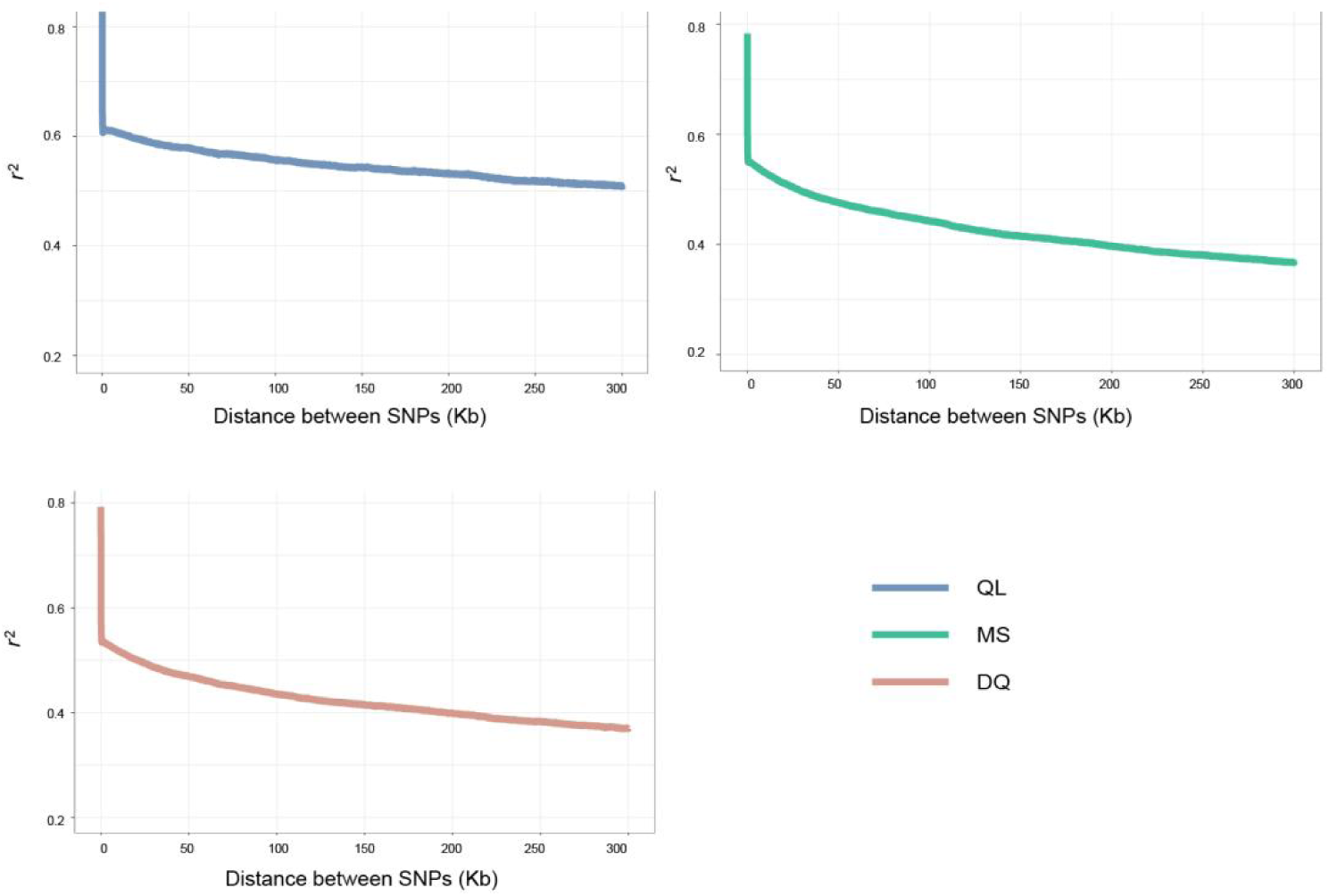
Decay of linkage disequilibrium (LD) with physical distance in three genetic groups of *Kingdonia uniflora*. Averages of pairwise linkage disequilibrium measures *r*^2^ are plotted for each bin of distances between pairs of SNPs. The displayed data are for the bins with pairs of SNPs separated by ≤300 kb

We used the ratio of nonsynonymous to synonymous polymorphisms (*π*_N_/*π*_S_) to assess the efficiency of purifying selection in *K. uniflora*. We defined a total of 24 genotypic groups according to multidimensional scaling (MDS) clustering (Fig. 5a). Among the 24 genotypic groups, genetic diversity at zero-fold degenerate (nonsynonymous) and four-fold degenerate (synonymous) sites (*π*_0_ and *π*_4_) is 0.000546 and 0.000992, respectively, resulting in *π*_0_/*π*_4_ (*π*_N_/*π*_S_)=0.55 (Fig. 5b).

**Fig. 5.**
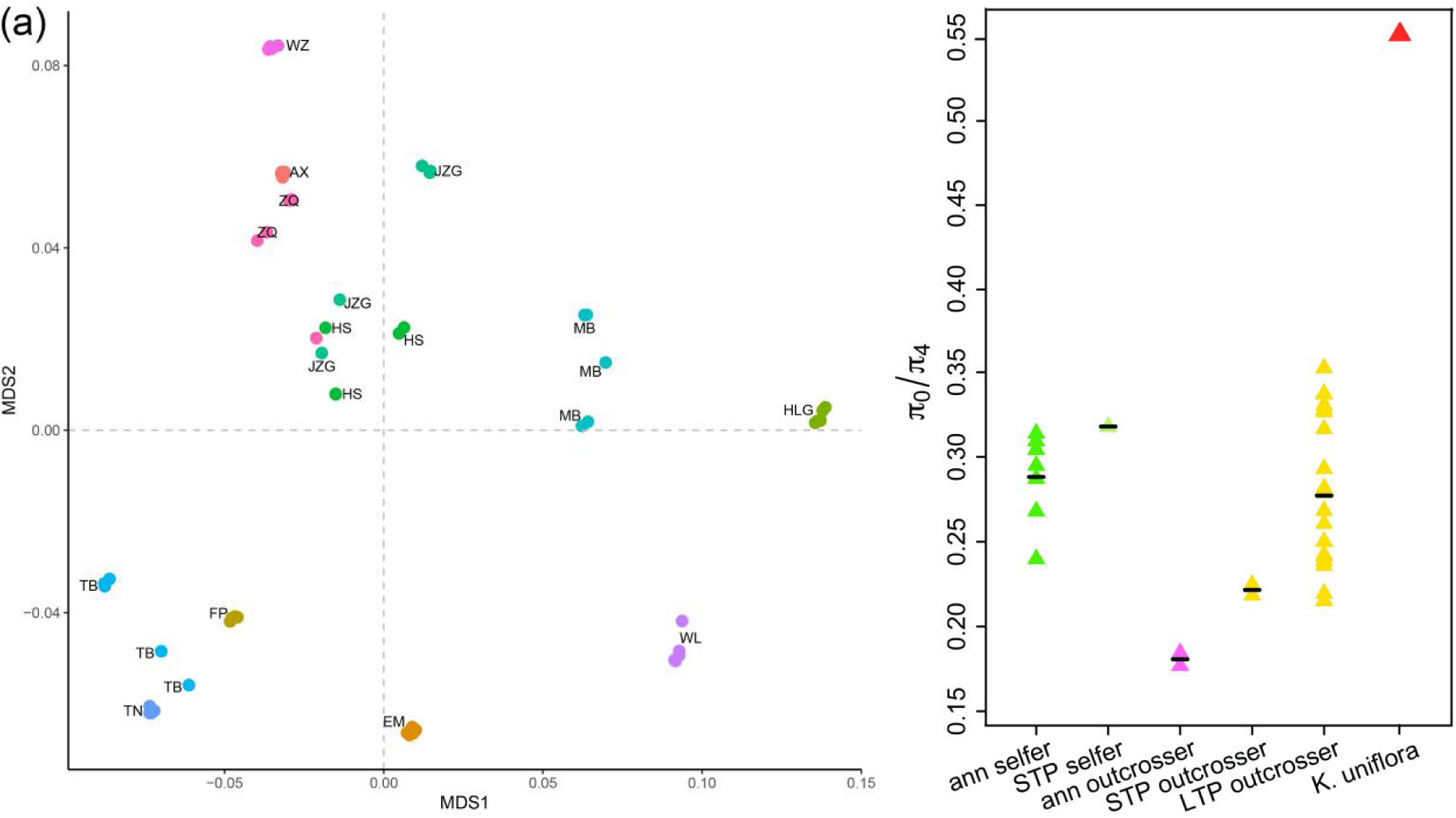
Calculation of *π*_N_/*π*_S_ in *K. uniflora*. (a) Multidimensional scaling analysis of identity-by-state pairwise distances between individuals. Individuals in the same population are coded with the same color. Individuals that come from the same population and cluster together are grouped into one genotypic group, and a total of 24 genotypic groups are identified. (b) *π*_0_/*π*_4_ difference between *K. uniflora* and other plants reported in Chen et al. (2017). The figure is modified based on Fig. 2B in Chen et al. (2017). STP, short-term perennial; LTP, long-term perennial.

Results from SplitsTree v4.13.1 (Huson and Bryant 2006) showed reticulations on the phylogeny of *K. uniflora*, which are signals of recombination (Fig. 6a). When all SNPs were concatenated into a single sequence, the RDP4 (Recombination Detection Program 4) (Martin et al. 2015) analysis identified 15 recombination events that were unevenly distributed among populations (Supplemental Table S4; Fig. 6b).

**Fig. 6.**
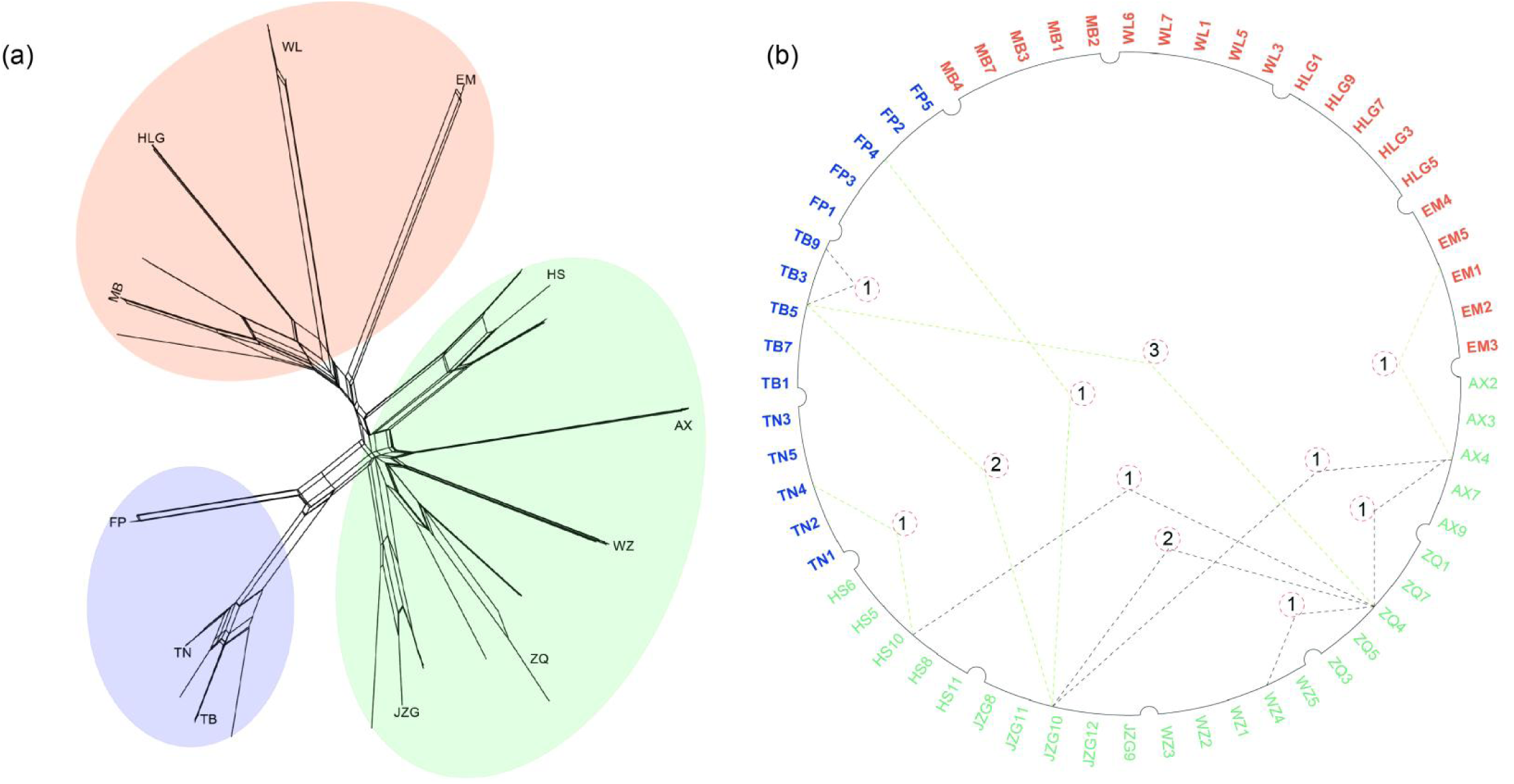
Recombination events detected in *Kingdonia uniflora* (a) Recombination events identified by SplitsTree phylogenetic networks using a total of 114,476 SNPs . The closed loops in the network shows recombination events. (b) Recombination events identified by RDP4. Black dotted lines indicate recombination between different individuals in the same genetic group; green dotted lines represent recombination between different individuals from different genetic groups; the number of corresponding DNA recombination events are indicated in circles.

### Inferring population demographic history and distributions

The model with the highest score found by simulations of demographic history using FSC2 (*fastsimcoal2*) (Excoffier et al. 2013) showed a divergence between the DQ and QL + MS groups 513,000 years ago and a divergence between the MS and QL groups 37,800 years ago (Table 3; Fig. 7a). The population sizes of DQ, MS and QL were estimated to be 98,800, 32,300 and 23,200, respectively (Table 3; Fig. 7a). The ancestral effective population size of the species (*N*_A1;_ Fig.7a) was estimated to be 29,000 (Table 3), slightly larger than that of QL, but smaller than those of DQ and MS (Table 3; Fig. 7a). The model also indicates asymmetric gene flow among the three genetic groups, with the rates of migration from QL to DQ and MS being higher than in the reverse direction. Between DQ and MS, the rate of migration from MS to DQ is higher than in the opposite direction (Table 3).

**Fig. 7.**
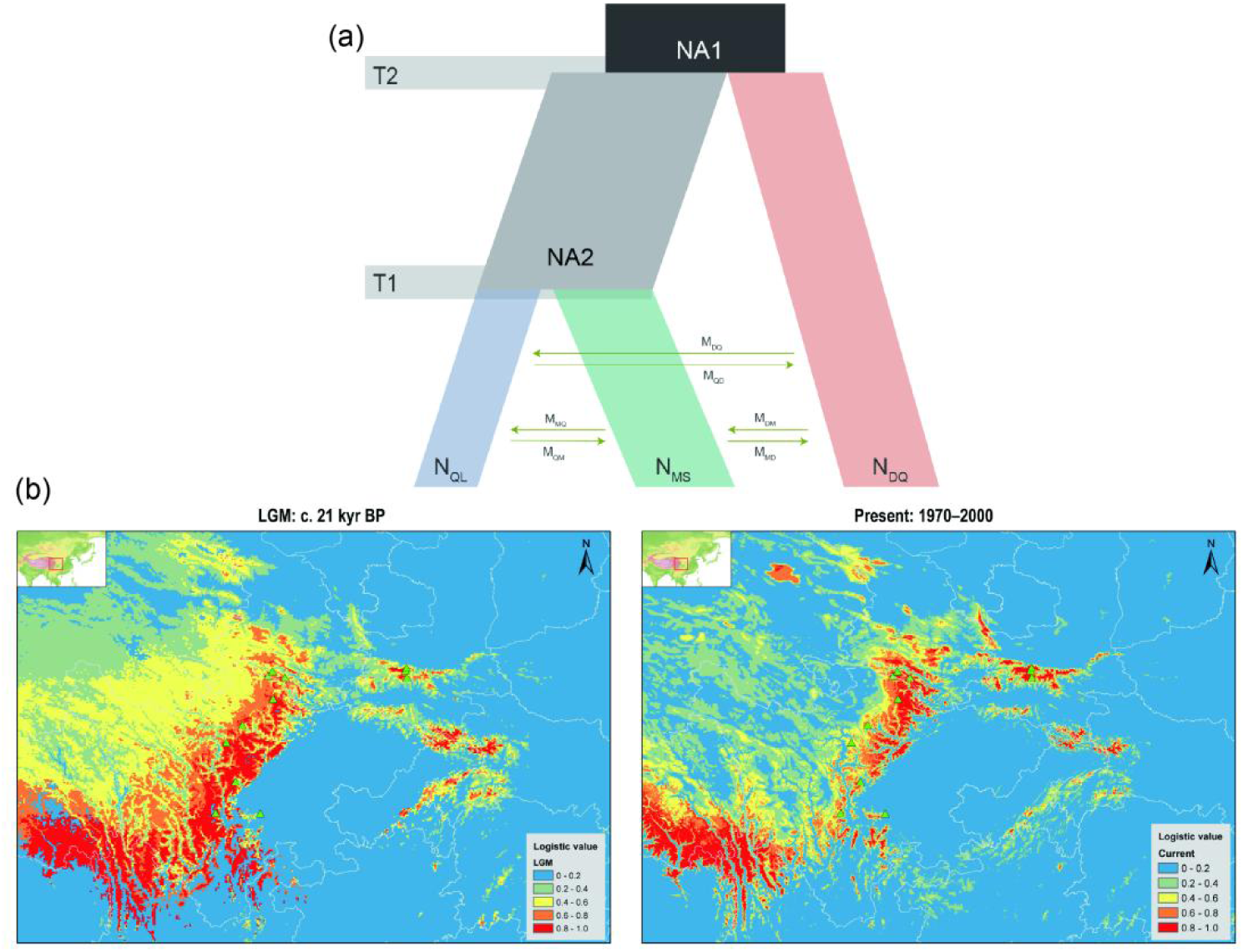
Inferred evolutionary history of *K. uniflora*. (a) Schematic of demographic scenarios modeled using FSC2, with the ancestral population shown in gray. Column width represents the relative effective population size. Arrows indicate gene flow between populations. (b) Predicted suitable distributions of *K. uniflora* at different historical periods based on species distribution modeling (SDM). Area color indicates probability (0-1) of suitable habitat for *K. uniflora*. LGM, last glacial maximum. The map image was derived from ArcGIS v10.3.

**Table 3.** Inferred demographic parameters for the best-fitting FSC2 model shown in Figure 7a, including 95% confidence intervals.

| Parameter | Point estimation | 95% confidence intervals |  |
| --- | --- | --- | --- |
|  |  | Lower bound | Upper bound |
| T<sub>1</sub> | 3.78E+04 | 2.65E+04 | 4.91E+04 |
| $T_2$ | 5.13E+05 | 4.60E+05 | 5.65E+05 |
| $N_{A1}$ | 2.90E+04 | 2.46E+04 | 3.33E+04 |
| $N_{A2}$ | 1.08E+05 | 1.00E+05 | 1.15E+05 |
| $N_{DQ}$ | 9.88E+04 | 6.58E+04 | 1.32E+05 |
| $N_{MS}$ | 3.23E+04 | 2.74E+04 | 3.71E+04 |
| $N_{QL}$ | 2.32E+04 | 3.55E+03 | 4.29E+04 |
| $M_{DM}$ | 2.30E-04 | 9.76E-05 | 3.63E-04 |
| $M_{DQ}$ | 1.53E-04 | 8.08E-05 | 2.26E-04 |
| $M_{MD}$ | 6.13E-04 | 3.26E-04 | 9.01E-04 |
| $M_{MQ}$ | 4.13E-04 | 1.58E-04 | 6.67E-04 |
| $M_{QD}$ | 1.08E-03 | 4.67E-04 | 1.69E-03 |
| $M_{QM}$ | 1.12E-03 | 6.84E-04 | 1.56E-03 |

Notes: Parameters included here comprise population size measures ( $N_{A1}$ , $N_{A2}$ , $N_{DQ}$ , $N_{MS}$ , and $N_{QL}$ , indicating ancestral populations, QL+MS, DQ, MS and QL, respectively), population divergence times ( $T_1$ and $T_2$ , years), migration per generation after diversification between QL and MS, QL and DQ, and DQ and MS.

Results from MAXENT ENM (ecological niche modeling) (Phillips and Dudik 2008) analyses showed a greater suitable habitat for *K. uniflora* in the LGM than at present, especially in the Hengduan Mountains region, and that there is more suitable habitat at present than is currently occupied (Fig. 7b).

### Assessing the correlation between genetic differentiation and environmental variables

Among the seven variables used for the GF (gradient forest) analysis, precipitation seasonality (bio15) and temperature seasonality (bio04) were identified as the two most important predictors of genetic variation. Mean temperatures of the warmest quarter (bio10), annual precipitation (bio12) and precipitation of the wettest month (bio13) were also of high importance (Fig. 8). The redundancy analysis (RDA) revealed a significant amount of genetic variation among populations associated with the seven important environmental variables (27.06%, *P* = 0.001). Each of the two axes explained a significant amount of variation (Axis 1: 25.80%, *P* = 0.001**; Axis 2: 17.03%, *P* = 0.001**; Fig. 8). Separate analyses of the seven environmental variables individually also found significant association with genetic variation (Table 4). Consistent with the GF analysis, temperature seasonality (bio04; 6.498%, P = 0.001**) and precipitation seasonality (bio15; 6.27%, P = 0.001**) were the two most important predictors (Table 4).

**Fig. 8.**
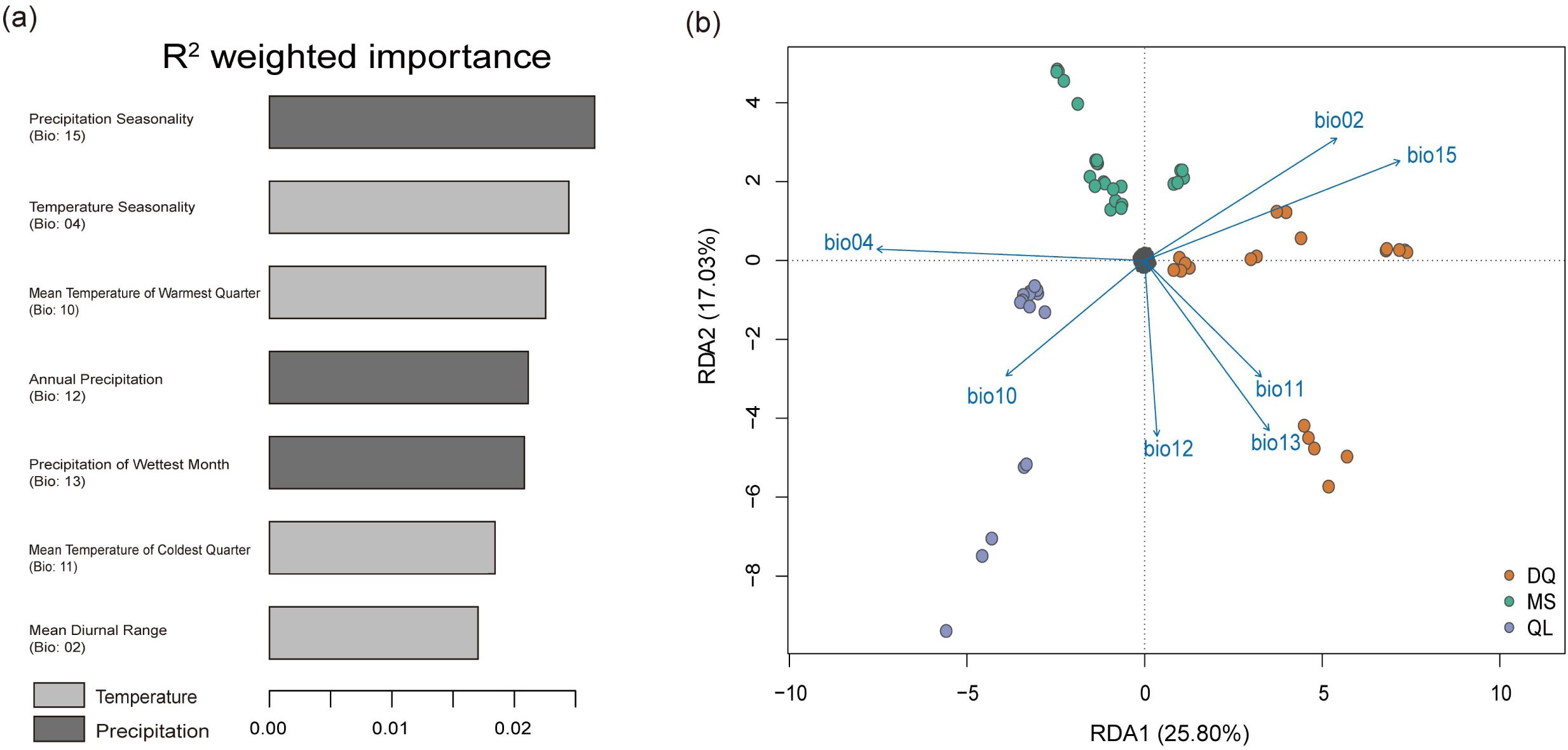
Correlation between genetic distance and environmental difference. (a) *R^2^-*weighted importance of environmental variables that explain genomic variation from GF analysis. (b) Redundancy analysis showing the relationship between the independent climate parameters and population structure of *K. uniflora*. Individuals are colored points and colors represent three groups (QL, MS, DQ). Small black points are SNPs.

**Table 4.**
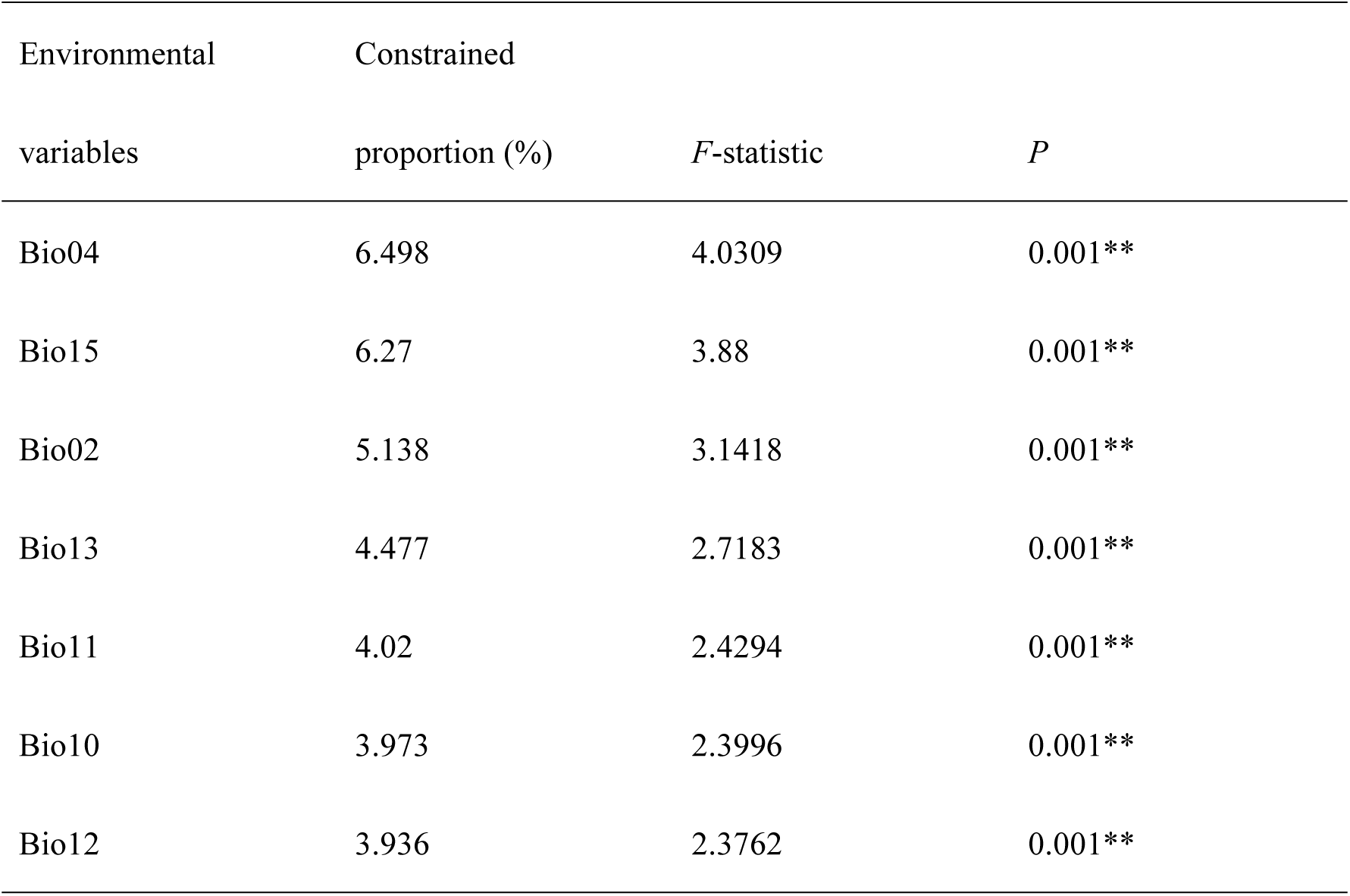
Redundancy analysis (RDA) results based on seven important environmental variables identified by GF analysis. **, *P* < 0.01.

### Detecting and annotating outlier loci

The average *F*_ST_ between populations was estimated as 0.24, but ranged from 0.05 to 0.76 (Fig. 9). BAYESCAN identified 7,917 outlier (*F*st values ranging from 0.47 to 0.76) sites out of the total 114,746 SNPs. Through comparisons with the reference genome, we found these outliers were located in 159 genes, of which 87 had BLAST hits in the Unified Protein Database, related to 15 biological processes (second-level gene ontology (GO) terms) (Fig. 10a). Gene set enrichment (GSE) analysis for all 87 genes detected five significantly over-represented terms (*p* < .05) associated with reproduction, including: “reproduction” (13 genes), “reproductive process” (13 genes), “reproductive developmental process” (11 genes), “fruit development” (6 genes), and “seed development” (6 genes) (Fig. 10b; Table 5).

**Fig. 9.**
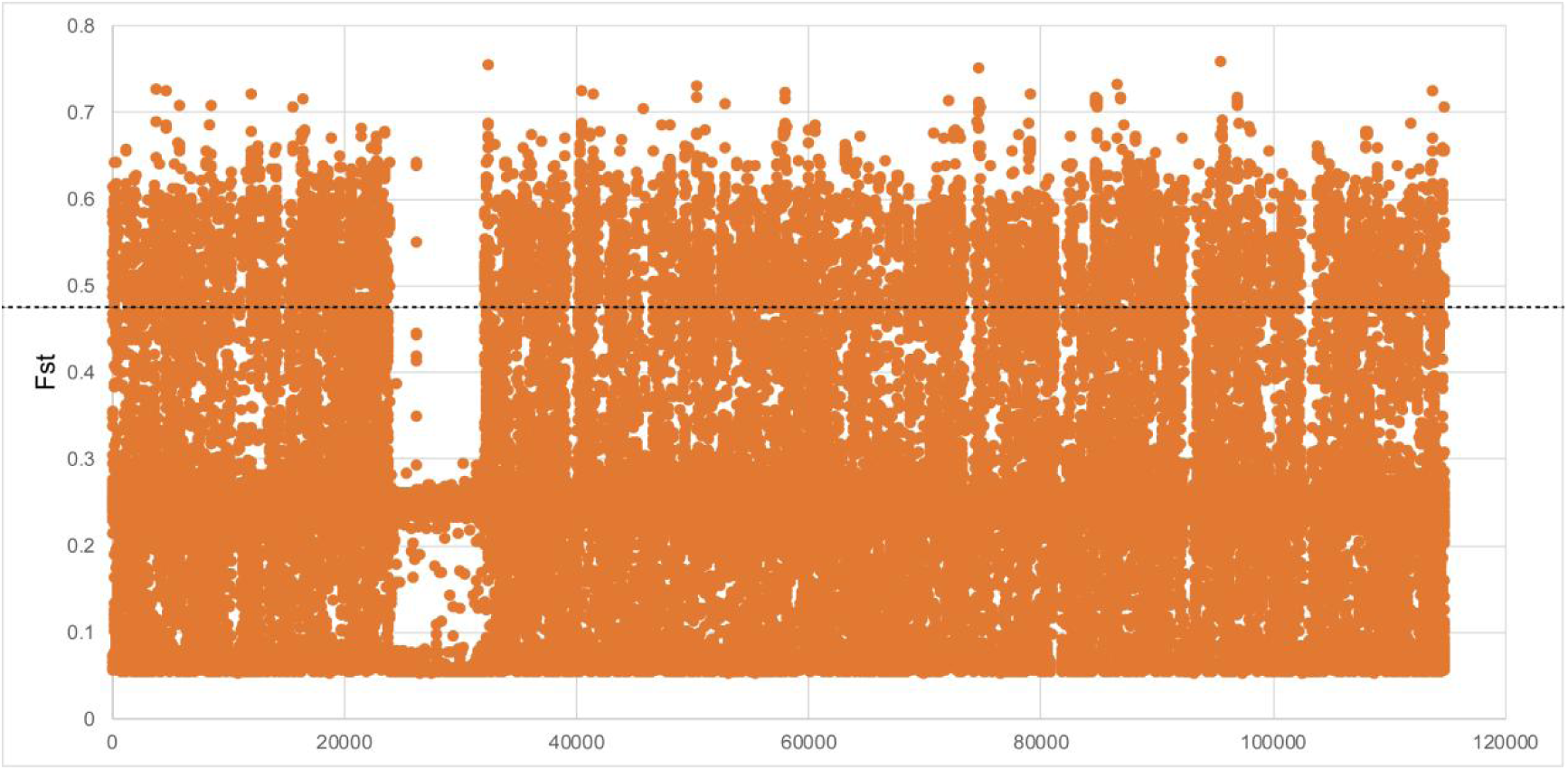
F_S_T values of 1, 114,746 SNPs among 60 *K. uniflora* individuals calculated by BAYESCAN. The x-axis represents the sequence number of SNPs. The black dash line indicates the cut-off line between identified outliers and non-outliers.

**Fig. 10.**
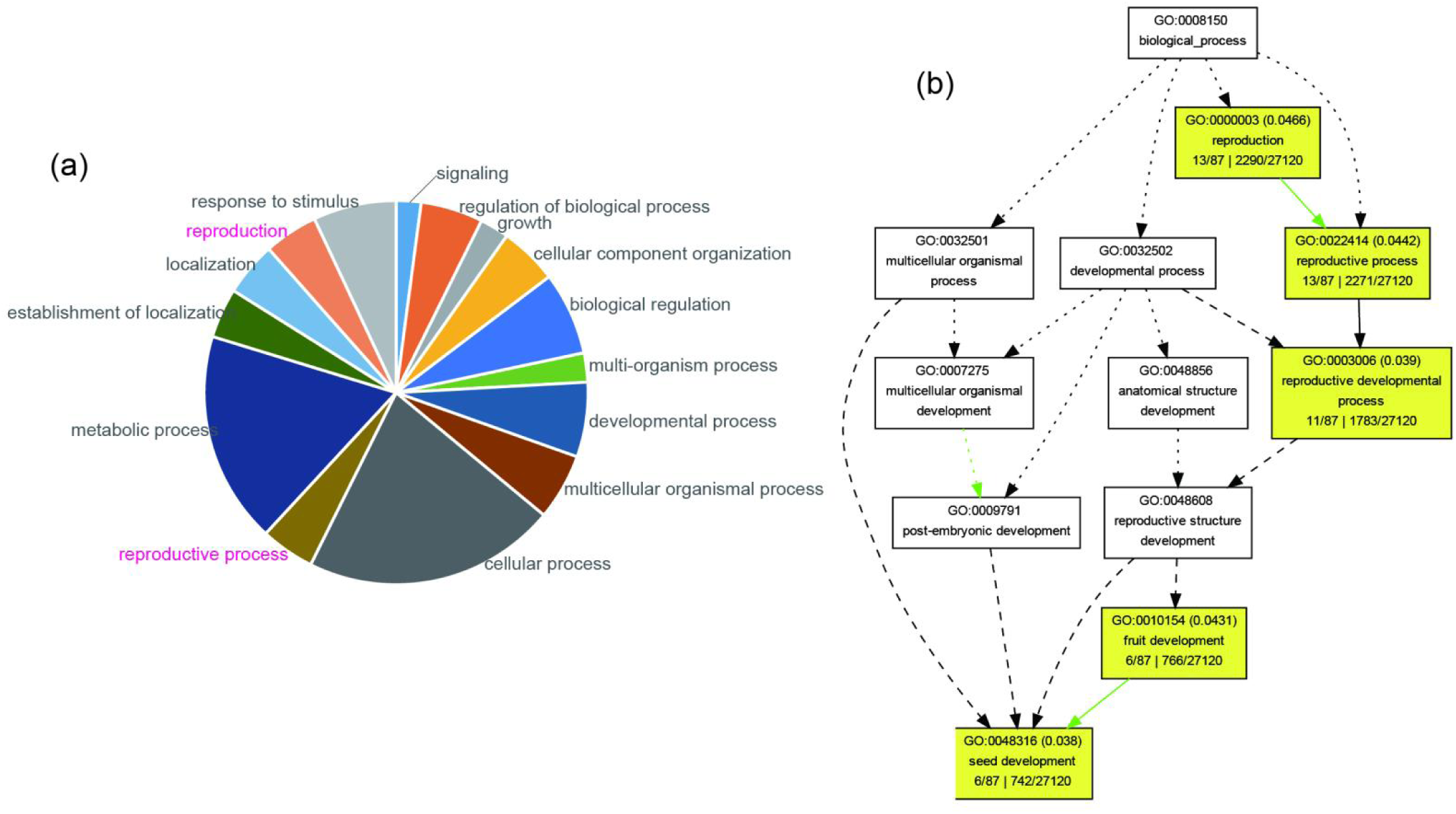
Gene set enrichment (GSE) analysis of genes containing outlier SNPs. (a) Pie chart of second-level gene ontology (GO) terms (biological process) of genes showing outlier *F*st values. (b) Graphical results showing enriched GO terms (second to sixth level; highlighted with yellow color) of genes containing outlier SNPs.

**Table 5.** The significantly over-represented GO terms of genes containing outlier SNPs in K. uniflora.

| GO term | Class | Description | Significant | Annotated | *p*-value |
| --- | --- | --- | --- | --- | --- |
| GO:0048316 | P | seed development | 6 | 742 | 0.038 |
| GO:0003006 | P | reproductive developmental process | 11 | 1783 | 0.039 |
| GO:0010154 | P | fruit development | 6 | 766 | 0.043 |
| GO:0022414 | P | reproductive process | 13 | 2271 | 0.044 |
| GO:0000003 | P | reproduction | 13 | 2290 | 0.047 |

## Discussion

### *Kingdonia uniflora* is characterized by high allelic heterozygosity, slow LD decay, reduced efficacy of purifying selection, and unseen sexual reproduction

In diploid asexuals, high levels of allelic divergence are expected to result from two factors: (1) long-term evolution under obligate asexuality, i.e., Meselson effect (Birky 1996; Mark Welch and Meselson 2000) and (2) transition to asexual reproduction via hybridization between sexual species (Jaron et al. 2021). The Meselson effect is usually considered to be a strong indicator of long-term evolution under obligate asexuality (Hartfield 2016; Brandt et al. 2021), yet this phenomenon has been shown to appear in young lineages of less than 100,000 years age (Pellino et al. 2013). Here we detected high allelic heterozygosity indicating obligate or high levels of asexuality in the flowering plant species *K. uniflora*: (1) an excess of observed individual heterozygosity over Hardy-Weinberg expectation as indicated by negative *F*_IS_ values (Fig. 2) (which is different from what is observed in the sister sexual species *C. agrestis* (mean *F*_IS_ = 0.02; Table 1 in Zhang et al. 2020) and (2) greater genetic divergence within individuals than that between populations revealed by the AMOVA analysis (Table 2). Theoretically, as mentioned above, both the Meselson effect and hybrid origin can explain high allelic heterozygosity in diploid asexuals. Hence high levels of allelic divergence detected in *K. uniflora* could be explained as accumulation of heterozygous variants caused by independent evolution of alleles after the transition to obligate or high levels of asexuality. Alternatively, *K. uniflora* could have switched to obligate or high level of asexuality via interspecific hybridization, as reported in asexual species of *Meloidogyne*, *Lineus* ribbon worms, and the *Ranunculus auricomus* complex (Lunt 2008; Pellino et al. 2013; Lunt et al. 2014; Ament-Velasquez et al. 2016). In the current study, although we cannot formally exclude a hybrid origin of asexuality, this is unlikely to be the case in *K. uniflora*. Different from the above hybrid origins of asexual species with multiple congeneric sister species, *Kingdonia* is a monotypic genus and has no fossil record that indicates the presence of parental species for hybridization. The closest related species, *C. agrestis*, is from a different genus and diverged from *K. uniflora* tens of millions of years ago (Sun et al. 2020; Zhang et al. 2020). The site frequency spectrum reveals the level of the heterozygous genotypes shared among lineages/populations greatly exceed those under Hardy-Weinberg equilibrium (Fig. 3), indicating high allelic heterozygosity occurred prior to lineage differentiation. Hence, in any event, both explanations (i.e., long-term asexuality and hybrid origin) provide support that *K. uniflora* has engaged in obligate or high levels of asexuality before lineage differentiation around 0.5 mya.

Asexual reproduction prevents free exchange of alleles among individuals and results in allele linkage disequilibrium (LD) and in extreme cases (obligate asexuality) may result in complete physical linkage of markers over the entire genome (Flint-Garcia 2003; Simko et al. 2006; Schurko et al. 2009). Various genetic processes in outcrossing can reduce LD and free random mating results in linkage equilibrium of alleles. Therefore, the level of LD in a species reflects the extent of inbreeding (non-random mating) or asexuality. Our data show that LD in *K. uniflora* has an average *r*^2^ up to 0.6, and the decay of LD with physical distance is extremely slow (Fig. 4). This pattern of LD decay is comparable to that seen in the highly self-fertilizing/asexual plant species, e.g., *Arabidopsis thaliana* (Nordborg et al. 2005; Kim et al. 2007), *Medicago truncatula* (Branca et al. 2011), and *Spirodela polyrhiza* (Ho et al. 2019), and completely different from outcrossing populations that often show rapid LD decay over several hundred bp (Foxe et al. 2009; Mackay et al. 2012; Ho et al. 2019). The LD decay pattern provides genetic evidence supporting *K. uniflora* as a species is undergoing reproduction by high asexuality.

Interference among loci caused by high linkage disequilibrium will decrease the efficiency of selection by preventing selection from acting individually on each locus (Gordo and Charlesworth 2001; Otto 2021). Such kinds of selective interference, e.g., a selective sweep, can result in a higher ratio of non-synonymous (selected) to synonymous (neutral) polymorphisms in asexual lineages (Ament-Velásquez et al. 2016; Hartfield 2016). The value of *π*_N_/*π*_S_ ratio (0.55) in *K. uniflora* is relatively higher than that (ranging from 0.20 to 0.35) observed in outcrossing plants (Fig. 5b), implying a higher rate of non-synonymous substitution caused by reduced efficacy of purifying selection in *K. uniflora*. Under high levels of asexuality, mutations are expected to be an important source of variation, with mutations typically occurring in a heterozygous state in asexual species due to independent evolution of alleles (as found in *K. uniflora* by the *F*_IS_ and SFS analyses). However, recessive mutations are not exposed to selection (Ament-Velásquez et al. 2016), which could be another reason why a higher rate of non-synonymous substitution is detected in *K. uniflora*.

Although high allele divergence, LD level and *π*_N_/*π*_S_ ratio are characterized by *K. uniflora*, low levels of sexual reproduction cannot be excluded. We detected recombination events in RDP4 analysis and genealogical network (Supplemental Table S4; Fig. 6), suggesting occurrence of sexual reproduction. The evidence supports *K. uniflora* as likely not an obligate asexual species. Although the detected recombination events could have been from mitotic recombination, due to the following evidence, we argue that occasional successful sexual reproduction exists in the species. First, the species still produces seeds, although no seedling has been observed in the field (Ren et al. 2003; Xu 2015). Second, high allele divergence has been shown to be compatible with low-rate sexual reproduction (Ceplitis 2003). Lastly, SVDQuartets showed a quarter of the quartets were incongruent with the species tree, indicating a portion of the SNPs did not diverge congruently with the rest of the SNP sites due to incomplete lineage sorting or recombination. Therefore, we hypothesize that the species likely engages, to an extent, in sexual reproduction to maintain genetic variation and slow down the speed of Muller’s ratchet, supporting the theory that recombination is necessary for long-term survival. In sum, our results indicate that *K. uniflora* is not a true obligate asexual species although the asexuality likely has evolved before 0.5 mya.

### Evolutionary history of *Kingdonia uniflora*

In asexual lineages, reduced efficacy of purifying selection caused by linked selection will lead to decreased fitness and a reduction in the effective population size *N*_e,_ (Nordborg and Donnelly 1997; Nordborg 2000; Ho et al. 2019). Populations with small *N*_e_ values usually show reduced capacity to respond to changing environmental pressures (Higgins and Lynch 2001; Siol et al. 2007). Currently, our knowledge of the evolutionary history of relict lineage with highly asexual reproduction, specifically the origins of genetic differentiation and demographic dynamics, has been extremely limited. In our study, results from the LEA and PAUP analyses indicate that populations within each of the three mountain systems, i.e., the Qinling Mountains, Minshan Mountains, and Daxue-Qionglai Mountains, belong to the same genetic cluster (Table 2; Fig. 1d). This observation suggests an important role of mountain isolation in shaping the population genetic structure of the species due to constraints in dispersal by distance (Fig. 1e) and/or abiotic environments, as evidenced by results from the GF and RDA analyses (Table 4; Fig. 8). The initial divergence of *K. uniflora* lineages was estimated to have occurred *c*. 0.51 million years ago (mya) (Table 3; Fig. 7a). The divergence time coincides with the occurrence of the Naynayxungla Glaciation (0.78-0.50 mya), one of the two largest glaciations on the Qinghai-Tibetan Plateau (QTP) (Zheng et al. 2002). We speculate that the climate caused by the Naynayxungla Glaciation triggered intraspecific differentiation in *K. uniflora* by shifting the distribution range, while the synergistic effects following geographic distance and environmental difference have further driven *K. uniflora* to eventually evolved into different genetic lineages.

Gene flow is an important way to weaken differentiation within species inhabiting montane regions, but is usually limited due to the aforementioned reasons. Gene flow in highly asexual species of flowering plants is especially limited due to the generally poor dispersal ability of propagules for asexual reproduction. Nevertheless, the FSC2 analysis support ancient gene flow between MS and both QL and DQ (Table 3; Fig. 7a). Our data from ecological niche modeling suggests gene flow likely occurred during the LGM when the species expanded its size and distribution to more suitable habitats as detailed below. Similar to many other cold adapted/tolerant species (e.g., Anderson et al. 2006; Tian et al. 2010; Opgenoorth et al. 2010; Gao et al. 2015), *K. uniflora* had a significant population expansion during the LGM (10,000-20,000 years ago) (Fig. 7b), well after the Naynayxungla Glaciation in the QTP or the divergence of the three lineages (Table 3; Fig. 7a). This provided opportunities for population admixture of the three previously (∼0.5 mya) diverged lineages of *K. uniflora*. Therefore, the gene flow likely have occurred recently during the LGM.

### Uneven distribution of sexual reproduction among *Kingdonia uniflora* populations is associated with differentiation in genes regulating seed development

Local adaptation involving diversifying selection is viewed as the best and longest manifestation of an evolutionary response to selection in nature (Kawecki and Ebert 2004; Lovell et al. 2014). Our data suggests such process occurred in *K. uniflora* in genes controlling seed development. The fact that field surveys and lab experiments failed to observe successful seed germination indicates failure or low rate of seed development and germination is likely to be the major factor that has led to non-evident (unseen/infrequent) sexual reproduction in *K. uniflora*. Our results showed that genes playing a strong role in divergence among *K. uniflora* populations are enriched for functions involved in seed development (Fig. 10; Table 5), suggesting differential selection of different alleles in different populations. An obvious divergence of DNA recombination frequency among lineages, e.g., extremely rare recombination events in DQ vs. relatively more frequent recombination events within QL and MS (Fig. 6b,7a) and between QL and MS, suggests different levels of sexual reproduction among and between population groups in different local environments. These together indicate the genetic divergence in seed development associated genes is likely to be the key factor of varied seed germination rate and sexual reproduction extent among *K. uniflora* populations. Although local adaptation might be constrained due to reduced efficacy of selection caused by genome-wide interference between loci (Jaron et al. 2021), our results show genetic differentiation in genes associated with seed development that, however, appear not to be random. The differences among populations in environmental variables explains a significant amount of genetic variation within *K. uniflora* (Fig. 8; Table 4), suggesting putative selection pressure on genes regulating seed development. This indicates the adaptive constraints posed by the high LD level in *K. uniflora* may be buffered by infrequent sexual reproduction. The differences in environmental variables, e.g., seasonal temperature and precipitation (Fig. 8), may directly affect seed germination rates and the corresponding genes among populations, further driving diversifying selection in shaping the diversity of genes regulating seed development. Overall, our results reveal that the infrequency or uneven distribution of sexual reproduction in *K. uniflora* is associated with genetic differentiation among populations.

## Materials and Methods

### Sample collection and resequencing

We carefully selected a total of 12 populations (Supplemental Table S5) to represent all known localities of *K. uniflora*. We collected fresh leaves of five individuals that were at least ten meters away from each other in each population to reduce the possibility that each sample was not a physiologically independent ramet. We dried the leaves in silica gel in the field, then stored them at -20°C before DNA extraction. Genomic DNA extraction, library construction, and amplification followed the protocols of Novogene (Beijing, China) (Supplementary Methods). All samples were sequenced using the Illumina HiSeq 4000 platform with a pair-end read length of 150 bp by Novogene (https://en.novogene.com/). Illumina raw reads were filtered by removing adapters and low-quality reads using Trim Galore v0.6.5 (https://www.bioinformatics.babraham.ac.uk/projects/trim_galore) with default options (Phred quality threshold 20; adapter auto-detection) (Krueger 2021)

### SNP calling and quality control

We mapped the filtered reads of each individual to the *K. uniflora* genome (Sun et al. 2020) using BWA-MEM v0.7.12-r1039 (Li 2013) with default parameters. We then converted sequence alignment/map (SAM) format files to BAM and sorted the BAM files using SAMtools v1.6 (Li et al. 2009) and conducted the following analyses in Genome Analysis Toolkit (GATK, v4.0) (Depristo et al. 2011). We first marked duplicate reads using MarkDuplicates. We then processed the data with AddOrReplaceReadGroups to replace all read groups in the INPUT file with a single new read group name and assign all reads to this read group in the OUTPUT BAM. To identify SNPs, we first conducted single-sample haplotype calling with HaplotypeCaller, and then identified Multi-sample SNPs using GenotypeGVCFs after merging the haplotype caller results from each sample using CombineGVCFs which aligned the haplotypes among samples. To obtain high quality SNPs, we filtered multi-sample SNPs using VariantFiltration with strict filter settings “QD < 2.0 || MQ < 40.0 || FS > 60.0 || SOR > 3.0 || MQRankSum < -12.5 || ReadPosRankSum < -8.0”. We further filtered the SNPs data to exclude monomorphic or triallelic variants, indels, and SNPs missing in any samples via VCFtools v0.1.16 (Danecek et al. 2011) for further analyses.

### Genetic diversity and population structure

To assess the population structure of *K. uniflora*, we performed the following analyses. We used Landscape and Ecological Association (LEA) (v3.3.2) R package (Frichot and François 2015) to determine the number of ancestral populations. LEA was developed for large genotypic matrices and does not rely on the genetic assumptions of the absence of genetic drift, Hardy-Weinberg or linkage equilibrium in ancestral populations (Pritchard et al. 2000). We also used a coalescent-based method to estimate a “species tree” based on the SNP data for comparison with results from LEA. Specifically, we used the SVDQuartets method (Chifman and Kubatko 2014) implemented in PAUP* v4.0a166 (Swofford, 2003; http://paup.phylosolutions.com/) with 100 bootstrap replicates and the quartet assembly method QFM to produce a species tree (Reaz et al. 2014).

We estimated nucleotide diversity (π) and genetic differentiation (*F*_ST_) between groups identified in the aforementioned analyses using VCFtools v0.1.16 (Danecek et al. 2011), and used Analysis of Molecular Variance (AMOVA) in Arlequin v3.5.2.2 (Excoffier and Lischer 2010) to estimate relative contributions of genetic variation from within and between groups. To determine if genetic differentiation is associated with geographic distance, we performed an IBD (isolation by geographic distance) analysis. We first estimated population-level genetic differentiation *F*_ST_ using the Weir and Cockerham method (Weir and Cockerham 1984) implemented in HIERFSTAT (Goudet 2005) in R v3.6.1 (R Core Team 2014). We then calculated genetic distance with the formula *F*_ST_/(1−*F*_ST_) and computed the pairwise geographic distance among 12 populations using GENALEX v6.5 (Peakall and Smouse 2012). We then tested the significance for the relationship between geographical distances and genetic distance among populations by conducting Mantel tests with ADE4 v1.7 (https://CRAN.R-project.org/package=ade4) using *mantel.rtest* with 9999 permutations.

### Detecting genomic signatures and signs of asexual and sexual reproduction

To understand the reproductive strategies, we first employed VCFtools v0.1.16 (Danecek et al. 2011) (option-het) to calculate *F*_IS_=1-*H*_obs_/*H*_exp_ for each individual, where *H*_obs_ and *H*_exp_ are the observed and expected heterozygosity, respectively. A negative *F*_IS_ value indicates an excess of observed individual heterozygosity. We then examined the distribution pattern of the excessive heterozygosity within and among the genetic groups identified in the aforementioned analyses by generating a site frequency spectrum (SFS) using Pop-Con with standard parameters (https://github.com/YoannAnselmetti/Pop-Con). Twelve randomly selected individuals that cover all populations were used for the SFS analysis.

We further examined other genetic consequences of asexual reproduction (i.e., linkage disequilibrium (LD) and reduced efficiency of purifying selection) and genetic evidence of unseen sexual reproduction (i.e., recombination). We calculated pairwise linkage disequilibrium (*r*^2^ value) and modeled the decline of LD with physical distance using PopLDdecay v3.40 (Zhang et al. 2019) with default settings. Within each contig we calculated LD between pairs of sites up to 300 kb and 1000 kb, respectively. In obligate asexuals, genome-wide LD between loci is expected, and the decline of LD between loci does not depend on their physical distance (Vakhrusheva et al. 2020).

To test if *K. uniflora* is characterized by reduced efficacy of purifying selection, we calculated the *π*_N_/*π*_S_ ratio of the species. For asexual species, individuals from the same population may be clones via descending from a common ancestor via only clonal reproduction, possibly over many generations (Ho et al. 2019). Therefore, prior to the calculation, we grouped individuals by conducting multidimensional scaling (MDS) analysis in plink v1.9 with the options -cluster, -mds-plot 2 eigvals and -allow-extrachr (Purcell et al. 2007) to group individuals that were genotypically highly similar into the same genotypic group within a population. We then randomly selected one individual from each genotypic group and generated a concatenated coding sequence for each selected individual based on SNP loci from coding regions using a custom script (Supplemental Methods). The sequence matrix comprising all above selected individuals was used for following nucleotide diversity calculation. Designated nucleotide diversity at 0-fold and 4-fold degenerate positions, *π*_N_ (*π*_0_) and *π*_S_ (*π*_4_), were calculated in MEGA (v10.1.6) (Kumar et al. 2018).

To assess if any unseen sexual reproduction may have occurred in *K. uniflora*, we reconstructed the relationship of individuals using SNP data with the NeighborNet method in SplitsTree v4.13.1 (Huson and Bryant 2006) with default settings. The method does not force a tree-like phylogeny in the analysis and can reveal phylogenetic networks resulting from recombination events. We further used Recombination Detection Program (RDP4) (Martin et al. 2015) to detect recombination events: “RDP4 implements a range of methods for both detection and characterization of recombination events that are evident within a sequence alignment without any prior user indication of a non-recombinant set of reference sequences” (Martin et al. 2015). We ran the analyses using six algorithms (RDP, BootScan, MaxChi, Chimera, GeneConv and SiScan) of recombination signal detection with Bonferroni correction and the highest acceptable *p* value 0.05. Recombination events that were identified by at least three of the six methods were considered.

### Inferring population demographic history and species distributions

To obtain a comprehensive view of the species’ evolutionary history, we first inferred the demographic history of *K. uniflora* using *fastsimcoal2* (FSC2; Excoffier et al. 2013) with the following details. We tested eight possible *N*_e_ models (Supplemental Fig. S5) to find the best model explaining our data for the genetic groups revealed by PAUP and LEA analyses. Then we estimated the composite likelihood of the observed data given a specified model using SFS (which was constructed here using easySFS.py instead of Pop-Con to generate a direct input file for FSC2) (https://github.com/isaacovercast/easySFS). Each model was run 20 times with 1,000,000 simulations for the calculation of the composite likelihood, and 40 expectation-conditional maximization (ECM) cycles. We compared the models based on the maximum likelihood value across 50 independent runs using the Akaike Information Criterion and Akaike’s weight of evidence and chose the model with the maximum Akaike’s weight value as the optimal model. Finally, we calculated confidence intervals of parameter estimates from 100 para-metric bootstrap replicates by simulating SFS from the maximum composite likelihood estimates and re-estimating parameters each time.

We also performed ecological niche modeling (ENM) to reconstruct the distribution range of the species in the present and past using MAXENT (Phillips and Dudik 2008) with population present occurrence data and climatic variables of the occurrence locations. Species’ presence occurrence data were compiled from the Chinese Virtual Herbarium (http://www.cvh.ac.cn) and our own field collections (Supplemental Table S5). We downloaded the climate layers of 19 bioclimatic variables (Supplemental Table S6) at a 2.5 arc minute resolution at present (average for the years 1970-2000) and during the last glacial maximum (LGM: *c.* 21 thousand years before present (kyr BP)) from WorldClim database website (http://www.worldclim.org/) (Fick and Hijmans 2017). To avoid multicollinearity, a Pearson correlation analysis was conducted to eliminate one of the variables in each pair with a correlation higher than 0.75, which resulted in seven climatic layers retained for analyses (Supplemental Table S6).

### Assessing the correlation between differentiation and environmental variables

We performed a gradient forest (GF) analysis implemented in GRADIENTFOREST v0.1 (http://gradientforest.r-forge.r-project.org/) to identify potential key environmental drivers of genomic variation in *K. uniflora*. The gradient forest method captures complex relationships between potentially correlated predictors (e.g., climatic variables) and multiple response variables (e.g., genetic variation), and provides the overall importance for each predictor weighted by *R^2^* (Ellis et al. 2012; Ma et al. 2020). The seven climatic layers retained in the ENM analysis were used as environmental predictors in the GF analysis for analyses of all SNPs. The analysis was run 1000 times to obtain the variability of *R^2^*, and the run with the highest overall performance (*R^2^*) for calculating weighted importance of predictors on response variables. To verify the results of GF analysis, we performed a redundancy analysis (RDA) to evaluate the associations between genetic variation and the seven environmental variables. We constrained the dependent variables (individuals) by the explanatory variables (environmental variables). The RDA analysis was performed using the rda function in VEGAN v2.5 (Oksanen et al. 2018; http://CRAN.R-project.org/package=vegan). The *anova.cca* function was used to calculate overall significance and significance of each climate variable using 9999 permutations.

### Detecting and annotating outlier loci

To detect potential loci under diversifying selection, we performed an overall *F*_ST_ outliers test in BAYESCAN v2.1 (Foll and Gaggiotti 2008) with default parameters. BAYESCAN identifies outliers using differences in allele frequencies between populations. A locus-specific component (*α*) was used to differ loci under-diversifying selection (*α*>0) from that under balancing or purifying selection. Significance is based on FDR-corrected *q*-values (<0.01). Loci under strong diversifying selection are those whose *F*st values are larger than expected from coalescent simulation of neutral evolution (Strand et al. 2012).

To determine what functions the genes containing SNP outliers may have, we annotated each of these genes using gene ontology (GO) terms with TBtools (Chen et al., 2018). We then used the Singular Enrichment Analysis (SEA) tool in agriGO v2.0 (Tian et al. 2017) to analyze gene enrichment and tested for statistical significance of gene enrichment using the chi-squared test.

## Data accessibility

*K. uniflora* resequencing reads have been deposited in the NCBI Short Read Archive (SRA) under accession SRA ------.

## Acknowledgments

This work was supported by the Program Foundation for the Backbone of Scientific Research by Wuhan Botanical Garden, Chinese Academy of Sciences (Y855241G01), the National Natural Science Foundation of China (U2003122), the Strategic Priority Research Program of Chinese Academy of Sciences (XDA20050203), and the National Natural Science Foundation of United States (DEB-1442161).

## Author contributions

HW, JX, and HS developed the idea and designed the experiment; YS, XZ, and HZ collected the leaf materials; YS, XZ, and AZ performed the statistical analyses; YS, JBL, and JX interpreted the results and wrote the manuscript. All authors read, edited and approved the final manuscript. YS, XZ, and AZ contributed equally to this work.

## References

Ament-Velasquez SL, Figuet E, Ballenghien M, Zattara EE, Norenburg JL, Fernandez-Alvarez FA, Bierne J, Bierne N, Galtier N. 2016. Population genomics of sexual and asexual lineages in fissiparous ribbon worms (Lineus, Nemertea): hybridization, polyploidy and the Meselson effect. Mol Ecol 25: 3356–3369. doi: 10.1111/mec.13717

Anderson LL, Hu SF, Nelson DM, Petit RJ, Paige KN. 2006. Ice-age endurance: DNA evidence of a white spruce refugium in Alaska. Proc Natl Acad Sci USA 13: 12447–12450. doi: 10.1073/pnas.0605310103

APG IV. 2016. An update of the angiosperm phylogeny group classification for the orders and families of flowering plants. Bot J Linn Soc 181: 1–20. doi: 10.1111/boj.12385

Balloux F, Lehmann L, de Meeûs T. 2003. The population genetics of clonal and partially clonal diploids. Genetics 164: 1635–1644. doi: 10.1093/genetics/164.4.1635

Beck JB, Alexander PJ, Allphin L, Al-Shehbaz IA, Rushworth C, Bailey CD, Windham MD. 2012. Does hybridization drive the transition to asexuality in diploid Boechera? Evolution 66: 985–995. doi: 10.1111/j.1558-5646.2011.01507.x.

Birky CW, Wolf C, Maughan H, Herbertson L, Henry E. 2005. Speciation and selection without sex. Hydrobiologia 546: 29–45. doi: 10.1007/s10750-005-4097-2

Birky CW. 1996. Heterozygosity, heteromorphy, and phylogenetic trees in asexual eukaryotes. Genetics 144: 427–437. doi: 10.1093/genetics/144.1.427

Branca A, Paape TD, Zhou P, Briskine R, Farmer AD, Mudge J, Bharti AK, Woodward JE, May GD, Gentzbittel L, et al. 2011. Whole-genome nucleotide diversity, recombination, and linkage disequilibrium in the model legume *Medicago truncatula*. Proc Natl Acad Sci USA 108: E864–E870. doi: 10.1073/pnas.1104032108

Brandt A, Tran Van P, Bluhm C, Anselmetti Y, Dumas Z, Figuet E, Francois CM, Galtier N, Heimburger B, Jaron KS, et al. 2021. Haplotype divergence supports long-term asexuality in the oribatid mite *Oppiella nova*. Proc Natl Acad Sci USA 118: e2101485118. doi: 10.1073/pnas.2101485118

Ceplitis A. 2003. Coalescence times and the Meselson effect in asexual eukaryotes. Genet Res 82: 183–190. doi: 10.1017/s0016672303006487

Chen C, Chen H, He YH, Xia R. 2018. TBtools, a Toolkit for Biologists integrating various biological data handling tools with a user-friendly interface. DOI: https://doi.org/10.1101/289660

Chen J, Glémin S, Lascoux M. 2017. Genetic diversity and the efficacy of purifying selection across plant and animal species. Mol Biol Evol 34: 1417–1428. doi:10.1093/molbev/msx088

Chifman J, Kubatko L. 2014. Quartet inference from SNP data under the coalescent model. Bioinformatics 30: 3317–3324. doi: 10.1093/bioinformatics/btu530

Corley LS, Blankenship JR, Moore AJ. 2001. Genetic variation and asexual reproduction in the facultatively parthenogenetic cockroach *Nauphoeta cinerea* : implications for the evolution of sex. J Evolution Biol 14: 68–74. doi: 10.1046/j.1420-9101.2001.00254.x

Danecek P, Auton A, Abecasis G, Albers CA, Banks E, DePristo MA, Handsaker RE, Lunter G, Marth GT, Sherry ST, et al. 2011. The variant call format and VCFtools. Bioinformatics 27: 2156–2158. doi: 10.1093/bioinformatics/btr330

de Meeûs T, Balloux F. 2004. Clonal reproduction and linkage disequilibrium in diploids: a simulation study. Infect Genet Evol 4: 345–351. doi: 10.1016/j.meegid.2004.05.002

DePristo MA, Banks E, Poplin R, Garimella KV, Maguire JR, Hartl C, Philippakis AA, del Angel G, Rivas MA, Hanna M, et al. 2011. A framework for variation discovery and genotyping using next-generation DNA sequencing data. Nat Genet 43: 491–498. doi: 10.1038/ng.806

Ellis N, Smith S, Pitcher C. 2012. Gradient forests: calculating importance gradients on physical predictors. Ecology 93: 156–168. doi: 10.1890/11-0252.1

Excoffier L, Dupanloup I, Huerta-Sanchez E, Sousa VC, Foll M. 2013. Robust demographic inference from genomic and SNP data. PLoS Genet 9: e1003905. doi: 10.1371/journal.pgen.1003905

Excoffier L, Lischer H. 2010. Arlequin suite ver 3.5: A new series of programs to perform population genetics analyses under Linux and Windows. Mol Ecol Resour 10: 564–567. doi: 10.1111/j.1755-0998.2010.02847.x

Felsenstein J. 1974. The evolutionary advantage of recombination. Genetics 78: 737–756. doi:10.1093/genetics/78.2.737

Fick S, Hijmans R. 2017. WorldClim 2: new 1-km spatial resolution climate surfaces for global land areas. Int J Climatol 37: 4302–4315. doi: 10.1002/joc.5086

Flint-Garcia SA, Thornsberry JM, Buckler ESt. 2003. Structure of linkage disequilibrium in plants. Annu Rev Plant Biol 54: 357–374. doi: 10.1146/annurev.arplant.54.031902.134907

Foll M, Gaggiotti O. 2008. A genome-scan method to identify selected loci appropriate for both dominant and codominant markers: a bayesian perspective. Genetics 180: 977–993. doi: 10.1534/genetics.108.092221

Foxe JP, Slotte T, Stahl EA, Neuffer B, Hurka H, Wright SI. 2009. Recent speciation associated with the evolution of selfing in Capsella. Proc Natl Acad Sci USA 106: 5241–5245. doi: 10.1073/pnas.0807679106

Frichot E, François O. 2015. LEA: AnRpackage for landscape and ecological association studies. Methods Ecol Evol 6: 925–929. doi: 10.1111/2041-210X.12382

Gao Y-D, Zhang Y, Gao X-F, Zhu Z-M. 2015. Pleistocene glaciations, demographic expansion and subsequent isolation promoted morphological heterogeneity: A phylogeographic study of the alpine *Rosa sericea* complex (Rosaceae). Sci Rep 5: 11698. doi: 10.1038/srep11698

Gladyshev E, Meselson M. 2008. Extreme resistance of bdelloid rotifers to ionizing radiation. Proc Natl Acad Sci USA 105: 5139–5144. doi: 10.1073/pnas.0800966105

Goudet J. 2005. HIERFSTAT, a package for R to compute and test hierarchical F-statistics. Mol Ecol Notes 5: 184–186. doi: 10.1111/j.1471-8286.2004.00828.x

Hartfield M. 2016. On the origin of asexual species by means of hybridization and drift. Mol Ecol 25: 3264–3265. doi: 10.1111/mec.13713

Heethoff M, Domes K, Laumann M, Maraun M, Norton RA, Scheu S. 2007. High genetic divergences indicate ancient separation of parthenogenetic lineages of the oribatid mite *Platynothrus peltifer* (Acari, Oribatida). J Evolution Biol 20: 392–402. doi: 10.1111/j.1420-9101.2006.01183.x

Henry L, Schwander T, Crespi BJ. 2012. Deleterious mutation accumulation in asexual Timema stick insects. Mol Biol Evol 29: 401–408. doi: 10.1093/molbev/msr237

Higgins K, Lynch M. 2001. Metapopulation extinction caused by mutation accumulation. Proc Natl Acad Sci USA 98: 2928–2933. doi: 10.1073/pnas.031358898

Ho EKH, Bartkowska M, Wright SI, Agrawal AF. 2019. Population genomics of the facultatively asexual duckweed *Spirodela polyrhiza*. New Phytol 224: 1361–1371. doi: 10.1111/nph.16056

Huson D, Bryant D. 2006. Application of phylogenetic networks in evolutionary studies. Mol Biol Evol 23: 254–267. doi: 10.1093/molbev/msj030

Jaron KS, Bast J, Nowell RW, Ranallo-Benavidez TR, Robinson-Rechavi M, Schwander T. 2021. Genomic features of parthenogenetic animals. J Hered 112: 19–33. doi:10.1093/jhered/esaa031

Kawecki TJ, Ebert D. 2004. Conceptual issues in local adaptation. Ecol Lett 7: 1225–1241. doi: 10.1111/j.1461-0248.2004.00684.x

Kim S, Plagnol V, Hu TT, Toomajian C, Clark RM, Ossowski S, Ecker JR, Weigel D, Nordborg M. 2007. Recombination and linkage disequilibrium in *Arabidopsis thaliana*. Nat Genet 39: 1151–1155. doi: 10.1038/ng2115

Kumar S, Stecher G, Li M, Knyaz C, Tamura K. 2018. MEGA X: molecular evolutionary genetics analysis across computing platforms. Mol Biol Evol 35: 1547–1549. doi: 10.1093/molbev/msy096

Laine VN, Sackton T, Meselson M. 2020. Sexual reproduction in bdelloid rotifers. bioRxiv doi: 10.1101/2020.08.06.239590v4

Lei Y, Ren L, Li Z, Ren Y. 2000. Studies on vegetative reproduction pattern of *Kingdonia uniflora*. Acta Botanica Boreali-Occidentalia Sinica, 20: 432–435.

Li H, Handsaker B, Wysoker A, Fennell T, Ruan J, Homer N, Marth G, Abecasis G, Durbin R, 1000 Genome Project Data Processing Subgroup. 2009. the sequence alignment/map format and SAMtools. Bioinformatics 25: 2078–2079. doi: 10.1093/bioinformatics/btp352

Li H. 2013. Aligning sequence reads, clone sequences and assembly contigs with BWA-MEM. arXiv doi:1303.3997

Li J, Zhang W, Li H. 2003. Research on distribution pattern of rare and endangered plant Kingdonia uniflora population. Journal of Northwest Forestry University 18: 38–40.

Li J, Zhao J, Li L. 2004. Property of sexual reproduction of *Kingdonia uniflora* population in Mountain Taibai. Journal of Northwest Sci-Tech University of Agriculture and Forestry 32: 89–92.

Lovell JT, Grogan K, Sharbel TF, McKay JK. 2014. Mating system and environmental variation drive patterns of adaptation in *Boechera spatifolia* (Brassicaceae). Mol Ecol 23: 4486–4497. doi: 10.1111/mec.12879

Lunt DH, Kumar S, Koutsovoulos G, Blaxter ML. 2014. The complex hybrid origins of the root knot nematodes revealed through comparative genomics. Peer J 2: e356. doi: 10.7717/peerj.356

Lunt DH. 2008. Genetic tests of ancient asexuality in root knot nematodes reveal recent hybrid origins. BMC Evol Biol 8: 194. doi: 10.1186/1471-2148-8-194

Ma S, Tian Y, Li J, Yu H, Cheng J, Sun P, Fu C, Liu Y, Watanabe Y. 2020. Climate variability patterns and their ecological effects on ecosystems in the Northwestern North Pacific. Front Mar Sci 7: 546882. doi: 10.3389/fmars.2020.546882

Mackay TF, Richards S, Stone EA, Barbadilla A, Ayroles JF, Zhu D, Casillas S, Han Y, Magwire MM, Cridland JM, et al. 2012. The *Drosophila melanogaster* genetic reference panel. Nature 482: 173–178. doi: 10.1038/nature10811

Mark Welch D, Meselson M. 2000. Evidence for the evolution of bdelloid rotifers without sexual reproduction or genetic exchange. Science 288: 1211–1215. doi: 10.1126/science.288.5469.1211

Martin DP, Murrell B, Golden M, Khoosal A, Muhire B. 2015. RDP4: Detection and analysis of recombination patterns in virus genomes. Virus Evol 1: vev003. doi: 10.1093/ve/vev003

Matzk F, Meister A, Schubert I. 2000. An efficient screen for reproductive pathways using mature seeds of monocots and dicots. Plant J 21: 97–108. doi: 10.1046/j.1365-313x.2000.00647.x

Maynard Smith J. 1978. The evolution of sex. Cambridge University Press, Cambridge (United Kingdom).

Mogie M. 1992. The evolution of asexual reproduction in plants. Chapman & Hall, London.

Muller HJ. 1964. The relation of recombination to mutational advance. Mutat Res-Fund Mol M 1: 2–9. doi: 10.1016/0027-5107(64)90047-8

Nei M, Gojobori T. 1986. Simple methods for estimating the numbers of synonymous and nonsynonymous nucleotide substitutions. Mol Biol Evol 3: 418–426. doi: 10.1093/oxfordjournals.molbev.a040410

Neiman M, Meirmans S, Meirmans PG. 2009. What can asexual lineage age tell us about the maintenance of sex? Ann NY Acad Sci 1168: 185–200. doi: 10.1111/j.1749-6632.2009.04572.x

Nordborg M, Donnelly P. 1997. The coalescent process with selfing. Genetics 146: 1185–1195. doi: 10.1093/genetics/146.3.1185

Nordborg M, Hu TT, Ishino Y, Jhaveri J, Toomajian C, Zheng H, Bakker E, Calabrese P, Gladstone J, Goyal R, et al. 2005. The pattern of polymorphism in *Arabidopsis thaliana*. PLoS Biol 3: e196. doi: 10.1371/journal.pbio.0030196

Nordborg M. 2000. Linkage disequilibrium, gene trees and selfing: an ancestral recombination graph with partial self-fertilization. Genetics 154: 923–929. doi: 10.1093/genetics/154.2.923

Normark B, Judson O, Moran N. 2003. Genomic signatures of ancient asexual lineages. Biol J Linn Soc 79: 69–84. doi: 10.1046/j.1095-8312.2003.00182.x

Normark BB, Moran NA. 2000. Testing for the accumulation of deleterious mutations in asexual eukaryote genomes using molecular sequences. J Nat Hist 34: 1719–1729. doi: 10.1080/00222930050122147

Opgenoorth L, Vendramin GG, Mao K, Miehe G, Miehe S, Liepelt S, Liu J, Ziegenhagen B. 2010. Tree endurance on the Tibetan Plateau marks the world’s highest known tree line of the Last Glacial Maximum. New Phytol 185: 332–342. doi: 10.1111/j.1469-8137.2009.03007.x

Otto SP. 2021. Selective Interference and the Evolution of Sex. J Hered 112: 9–18. doi: 10.1093/jhered/esaa026

Peakall R, Smouse P. 2012. GenAlEx 6.5: genetic analysis in Excel. Population genetic software for teaching and research–an update. Bioinformatics 28: 2537–2539. doi: 10.1093/bioinformatics/bts460

Pellino M, Hojsgaard D, Schmutzer T, Scholz U, Horandl E, Vogel H, Sharbel TF. 2013. Asexual genome evolution in the apomictic *Ranunculus auricomus* complex: examining the effects of hybridization and mutation accumulation. Mol Ecol 22: 5908–5921. doi: 10.1111/mec.12533

Pettersson ME, Berg OG. 2006. Muller’s ratchet in symbiont populations. Genetica 130: 199. doi: 10.1007/s10709-006-9007-7

Phillips S, Dudik M. 2008. Modeling of species distributions with Maxent: new extensions and a comprehensive evaluation. Ecography 31: 161–175. doi: 10.1111/j.0906-7590.2008.5203.x

Pritchard J, Stephens M, Donnelly P. 2000. Inference of population structure using multilocus genotype data. Genetics 155: 945–959. doi: 10.3410/f.1015548.197423

Purcell S, Neale B, Todd-Brown K, Thomas L, Ferreira MA, Bender D, Maller J, Sklar P, de Bakker PI, Daly MJ, Sham PC. 2007. PLINK: a tool set for whole-genome association and population-based linkage analyses. Am Journal Hum Genet 81: 559–575. doi: 10.1086/519795

R Core Team. 2014. R version 3.6.1: A language and environment for statistical computing. In *R Foundation for Statistical Computing*. Vienna, Austria.

Ruiz-Sanchez E, Rodriguez-Gomez F, Sosa V. 2012. Refugia and geographic barriers of populations of the desert poppy, *Hunnemannia fumariifoli*a (Papaveraceae). Organisms Diversity & Evolution 12:133–143. doi: 10.1007/s13127-012-0089-z

Reaz R, Bayzid MS, Rahman MS. 2014. Accurate Phylogenetic Tree Reconstruction from Quartets: A Heuristic Approach. PLoS ONE 9: e104008. doi:10.1371/journal.pone.0104008

Ren Y, Wang ML, Hu ZH. 1998. Kingdonia, embryology and its systematic significance. Acta Phytotaxonomica Sinica 36: 423–427.

Ren Y, Li Z, Lei Y. 2003. Achene and seed abortion contribute to the rarity of *Kingdonia uniflora*. Isr J Plant Sci 51: 39–44. doi: 10.1560/y466-dtln-5qf2-d4yh

Schön I, Rossetti G, Martens K. 2009. Darwinulid ostracods: ancient asexual scandals or scandalous gossip? Lost Sex 217–240. doi: 10.1007/978-90-481-2770-2_11

Schurko AM, Neiman M, Logsdon JM, Jr.. 2009. Signs of sex: what we know and how we know it. Trends Ecol Evol 24: 208–217. doi: 10.1016/j.tree.2008.11.010

Schwander T. 2016. Evolution: the end of an ancient asexual scandal. Curr Biol 26: R233–235. doi: 10.1016/j.cub.2016.01.034

Signorovitch A, Hur J, Gladyshev E, Meselson M. 2015. Allele sharing and evidence for sexuality in a mitochondrial clade of bdelloid rotifers. Genetics 200: 581–590. doi: 10.1534/genetics.115.176719

Simko I, Haynes KG, Jones RW. 2006. Assessment of linkage disequilibrium in potato genome with single nucleotide polymorphism markers. Genetics 173: 2237–2245. doi: 10.1534/genetics.106.060905

Siol M, Bonnin I, Olivieri I, Prosperi JM, Ronfort J. 2007. Effective population size associated with self-fertilization: lessons from temporal changes in allele frequencies in the selfing annual *Medicago truncatula*. J Evolution Biol 20: 2349–2360. doi: 10.1111/j.1420-9101.2007.01409.x

Strand AE, Williams LM, Oleksiak MF, Sotka EE. 2012. Can diversifying selection be distinguished from history in geographic clines? A population genomic study of killifish (Fundulus heteroclitus). PLoS One 7: e45138. doi: 10.1371/journal.pone.0045138

Sun Y, Deng T, Zhang A, Moore MJ, Landis JB, Lin N, Zhang H, Zhang X, Huang J, Zhang X, Sun H, et al. 2020. Genome Sequencing of the Endangered *Kingdonia uniflora* (Circaeasteraceae, Ranunculales) Reveals Potential Mechanisms of Evolutionary Specialization. iScience 23: 101124. doi: 10.1186/s12864-017-3956-3

Tian S, Lopez-Pujol J, Wang HW, Ge S, Zhang ZY. 2010. Molecular evidence for glacial expansion and interglacial retreat during Quaternary climatic changes in a montane temperate pine (*Pinus kwangtungensis* Chun ex Tsiang) in southern China. Plant Syst Evol 284: 219–229. doi: 10.1007/s00606-009-0246-9

Tian T, Liu Y, Yan H, You Q, Yi X, Du Z, Su Z. 2017. agrigo v2.0: A GO analysis toolkit for the agricultural community, 2017 update. Nucleic Acids Res 45: W122–W129. doi: 10.1093/nar/gkx382

Vakhrusheva OA, Mnatsakanova EA, Galimov YR, Neretina TV, Gerasimov ES, Naumenko SA, Ozerova SG, Zalevsky AO, Yushenova IA, Rodriguez F, et al. 2020. Genomic signatures of recombination in a natural population of the bdelloid rotifer *Adineta vaga*. Nat Commun 11: 6421. doi: 10.1038/s41467-020-19614-y

Weir B, Cockerham C. 1984. Estimating F-statistics for the analysis of population structure. Evolution 38: 1358–1370. doi: 10.1111/j.1558-5646.1984.tb05657.x

Xu J, Cao B, Bai C. 2015. Prediction of potential suitable distribution of endangered plant *Kingdonia uniflora* in China with MaxEnt. Chinese Journal of Ecology 34: 3354–3359.

Zhang C, Dong SS, Xu JY, He WM, Yang TL. 2019. PopLDdecay: a fast and effective tool for linkage disequilibrium decay analysis based on variant call format files. Bioinformatics 35: 1786–1788. doi: 10.1093/bioinformatics/bty875

Zhang X, Sun Y, Landis JB, Zhang J, Yang L, Lin N, Zhang H, Guo R, Li L, Zhang Y, et al. 2020. Genomic insights into adaptation to heterogeneous environments for the ancient relictual *Circaeaster agrestis* (Circaeasteraceae, Ranunculales). New Phytol 228: 285–301. doi: 10.1111/nph.16669

Zheng B, Xu Q, Shen Y. 2002. The relationship between climate change and Quaternary glacial cycles on the Qinghai-Tibetan Plateau: review and speculation. Quatern Int 97: 93–101. doi: 10.1016/S1040-6182(02)00054-X

Zimmer C. 2009. Origins. On the origin of sexual reproduction. Science 324: 1254–1256. doi: 10.1126/science.324_1254

